# Interplay between persistent activity and activity-silent dynamics in prefrontal cortex during working memory

**DOI:** 10.1101/763938

**Authors:** J Barbosa, H Stein, R Martinez, A Galan, K Adam, S Li, J Valls-Solé, C Constantinidis, A Compte

## Abstract

Persistent neuronal spiking has long been considered the mechanism underlying working memory, but recent proposals argue for alternative, “activity-silent” substrates for memory. Using monkey and human electrophysiology, we show here that attractor dynamics that control neural spiking during mnemonic periods interact with activity-silent mechanisms in PFC. This interaction allows memory reactivation, which enhance serial biases in spatial working memory. Stimulus information was not decodable between trials, but remained present in activity-silent traces inferred from spiking synchrony in PFC. Just prior to the new stimulus, this latent trace was reignited into activity that recapitulated the previous stimulus representation. Importantly, the reactivation strength correlated with the strength of serial biases in both monkeys and humans, as predicted by a computational model integrating activity-based and activity-silent mechanisms. Finally, single-pulse TMS applied to human prefrontal cortex prior to trial start enhanced serial biases, demonstrating the causal role of prefrontal reactivations in determining working memory behavior.

## Introduction

The mechanisms by which information is maintained in working memory are still not fully understood. Ample evidence supports a role of sustained neural activity in prefrontal ^1–3^ and other cortices ^4, 5^, possibly supported by attractor dynamics in recurrently connected circuits ^6–9^. However, recent studies have argued that memories may be maintained without persistent firing rate tuning during memory periods ^10^. This “activity-silent” memory can be mediated by short-term synaptic plasticity ^11, 12^, and possibly also by other activity-dependent intrinsic mechanisms with a long time constant^13–15^ that could allow reactivation of memories from a latent storage. This computational proposal has received support in neuroimaging studies: in some working memory tasks, even if memory performance is good, stimulus information cannot be retrieved from neural recordings during delay, but later robustly reappears ^16^ during comparison or response periods (but see ref. ^17^)

The apparent incompatibility between activity-based and activity-silent memory maintenance has led to viewing them as exclusive alternatives ^10^. However, modeling studies that have successfully implemented activity-silent conditions invariably require the network to be configured close to the attractor regime ^11, 18^, a plausible mechanism for persistent activity. The attractor non-linearity is necessary to increase the contrast of the fading subthreshold signal for successful memory reactivation. At the same time, mechanisms used for activity-silent memory may play a supportive role for persistent activity in attractor networks ^13, 19–21^, albeit with the cost of serial biases ^13, 22^. Serial biases in spatial working memory tasks denote small but systematic biases in reporting the location memorized in the current trial slightly attracted to nearby locations memorized in the previous trial ^23–26^. That these apparently non-adaptive serial biases reflect the direct interaction of persistent activity and activity-silent mechanisms is a theoretically appealing hypothesis that still lacks experimental support ^13, 22, 27^.

Both attractor dynamics ^24, 28^ and activity-silent ^13, 22, 27^ mechanisms have been proposed to carry stimulus-selective information from one trial to the next to effect serial biases. However, dependencies of serial biases with delay and ITI durations ^24–26^, which demonstrate their relationship with memory maintenance, are largely consistent with activity-silent, and not activity-based mechanisms ^13, 22, 27^. Here, we sought to specify the interaction of activity-based and activity-silent prefrontal cortex (PFC) mechanisms in supporting serial biases while subjects performed a spatial working memory task that engages attractor dynamics in prefrontal cortex ^6^. We compared the encoding properties of brain activity in delay and inter-trial intervals (ITI) to identify the mechanistic basis of the memory trace that spans consecutive trials. We used behavioral and electrophysiological data collected in monkeys and humans, with prefrontal multiple-unit recordings in monkeys, and scalp electroencephalography (EEG) in humans. Between successive persistent activity mnemonic codes, we found an activity-silent code in PFC that carried stimulus information through inter-trial periods. In addition, we found correlational and causal evidence that fixation-period PFC reactivation from this activity-silent trace enhances attractive serial biases. These findings underscore the behavioral relevance of the dynamic interplay between attractor and subthreshold network dynamics in PFC and reconcile these seemingly conflicting mechanisms: their interplay could be the basis of closely associated memory storage processes operating at different time scales, possibly serving different behavioral purposes ^29, 30^.

## Results

We trained four rhesus monkeys to perform an oculomotor delayed response task (ODR). The task consisted of remembering spatial locations at fixed eccentricity while maintaining fixation during a delay period of 3 s (Fig. 1a, Methods). The extinction of the fixation cue triggered the monkey to execute a saccade towards the remembered location and marked the beginning of a fixed inter-trial interval (ITI) of 3.1 s, lasting until the appearance of the new trial’s stimulus cue (Fig. 1b). In addition, we tested 35 human participants in variations of the task performed by the monkeys (Methods). In all cases, we recorded the reported location and computed behavioral errors as angular distances to corresponding target locations. Following the methods described in previous studies ^23^, we analyzed the dependence of the current-trial error on relative previous trial location. Both monkeys and humans showed biased reports relative to previously remembered locations. These biases were attractive for short distances between previous-trial and current-trial locations, and repulsive for large previous-current distances (Fig. 1a, 2a). Our primary goal was to test the hypothesis that activity-silent and persistent activity working memory mechanisms interact to produce serial dependence effects. To this end, we investigated electrophysiological measurements in the ITI, including periods from the response to the subsequent fixation period.

**Figure 1.**
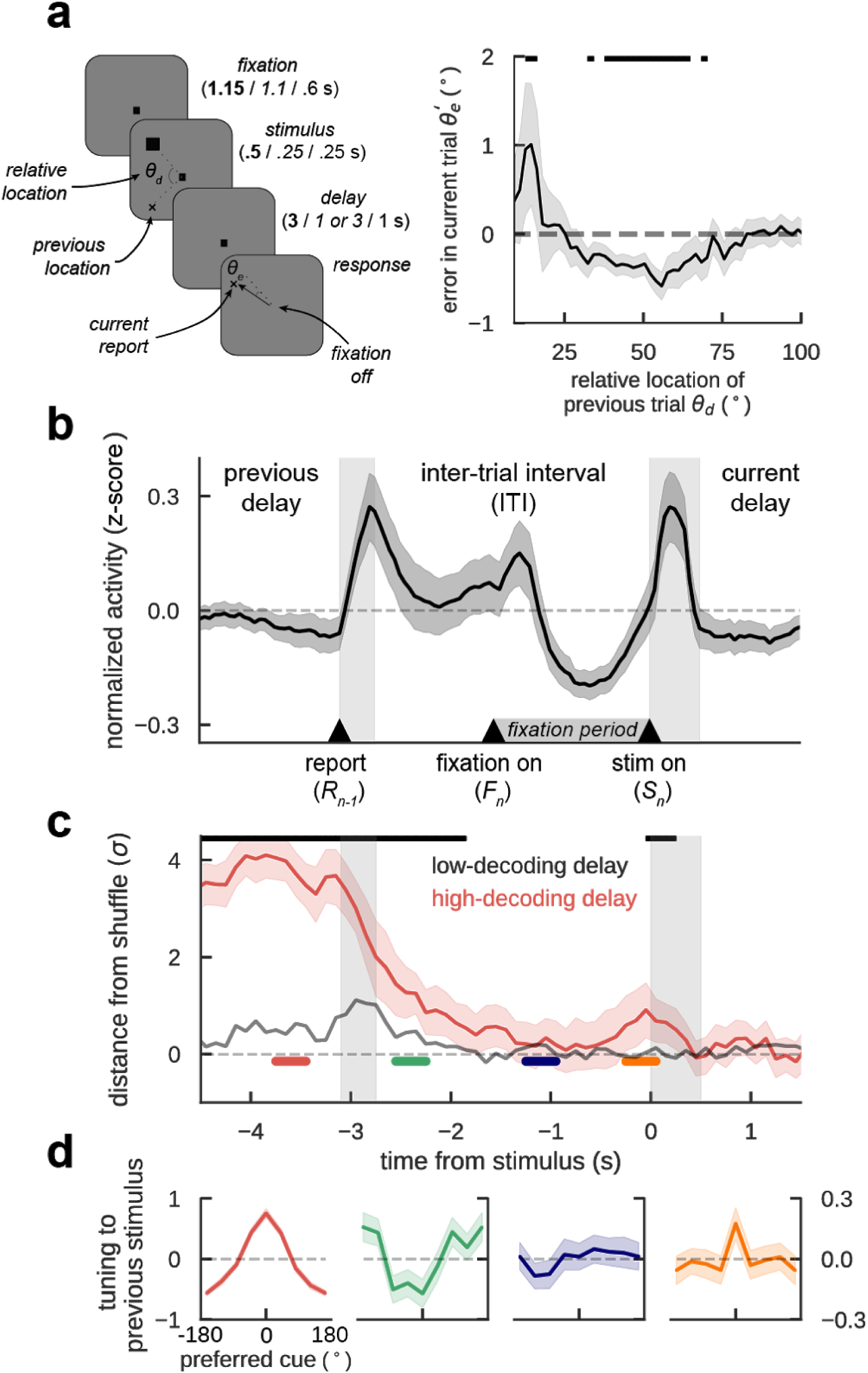
Previous-trial stimulus code reactivates prior to the forthcoming stimulus. **a)** General task design (**monkeys** / *EEG* / TMS) and serial bias for 4 monkeys. Trials with clockwise previous reports relative to the current stimulus were collapsed into clockwise trials (*folded errors*, Methods). Positive (negative) values indicate response attraction (repulsion) toward previous stimuli presented at that relative distance from the current stimulus. Shading indicates bootstrapped ±s.e.m. Black solid bars represent p<0.05 (permutation test). **b)** Averaged, normalized firing rate of n=206 neurons during the ITI (spike counts of 300-ms causal square kernel, z-scored in interval [-4.5 s, 1.5 s]). Gray bars mark response and stimulus cue periods. **c)** Decoding accuracy of previous-trial stimulus, computed as the distance from the mean of decoding accuracy in shuffled surrogates, in units of their standard deviation σ (Methods), averaged over ensembles with strong (red) and weak (gray) decoding in delay (Methods). Aligned with anticipatory ramping in late fixation (panel b), previous-trial stimulus code reappears, specifically in ensembles with better delay code (Supplementary Fig. 1). Black bars mark timepoints for which decoding accuracy 99.5% C.I. is above zero. **d)** Tuning to previous-trial stimulus, aligning responses to preferred cue as defined in delay, and computed in different trial epochs (color-coded in b, bootstrap-test at preferred location: p=0.015, c.i.=[-0.3,-0.03], Cohen’s d=-0.17 (green), p=0.865, c.i.=[-0.12,14], Cohen’s d=0.15 (blue), p=0.025, c.i.=[0.024,0.33], Cohen’s d=0.012 (orange), n=206 neurons, shading depicts ±s.e.m.). Unless stated otherwise, in all panels error-shading marks 95% C.I.

**Figure 2.**
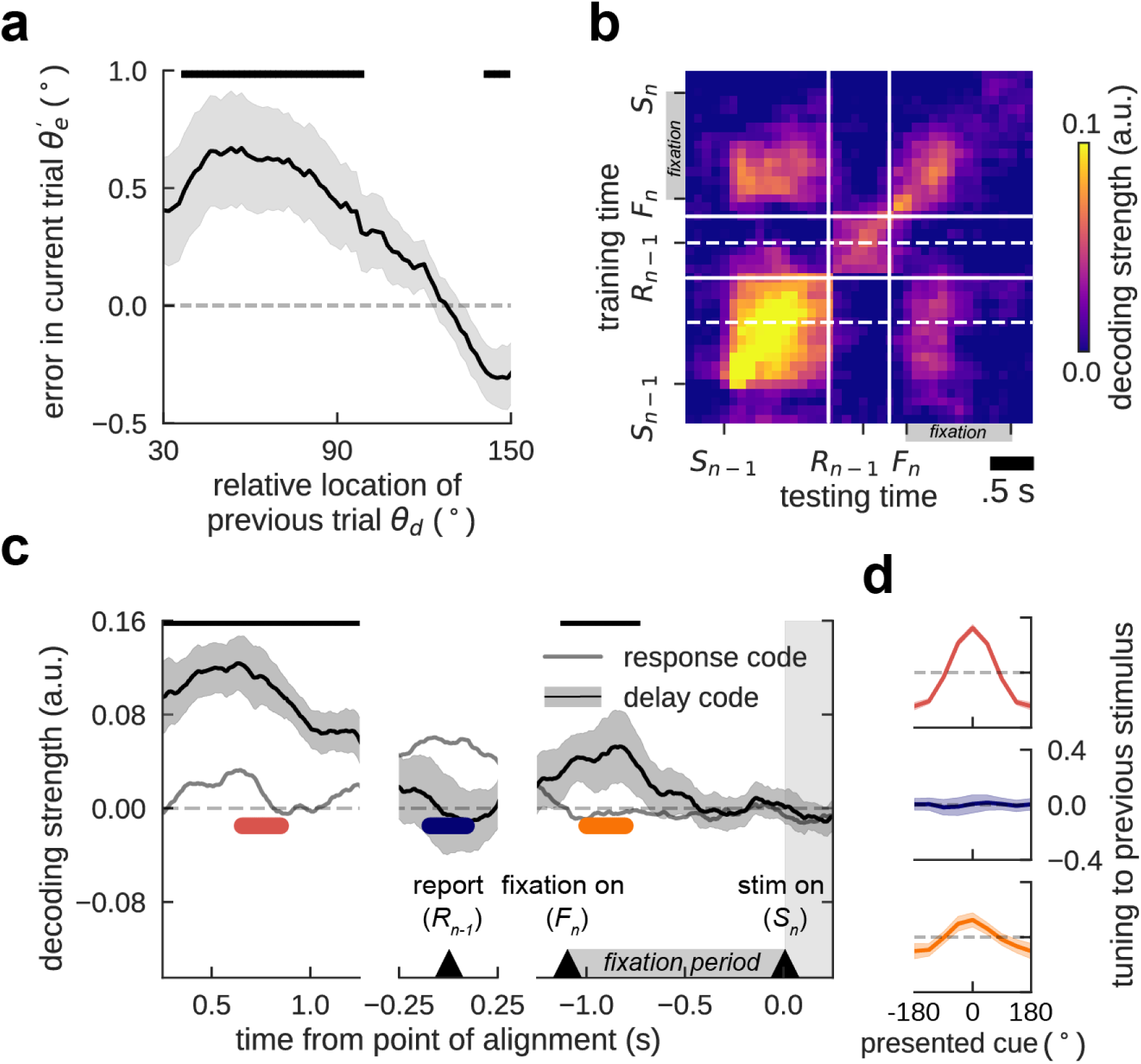
In human EEG, the delay code also reactivates in fixation. **a)** Folded serial bias (Methods) for 15 human subjects. Shading shows ±s.e.m. **b)** Temporal generalization of previous-stimulus code for all combinations of training and testing times (independent windows of 50 samples, ∼ 97.77ms) from previous trial stimulus onset (*S_n_*_−1_) and response (*R_n_*_−1_), to current trial fixation onset (*F_n_*) and stimulus onset (*S_n_*). Solid white lines mark the discontinuity of EEG fragments aligned to *S_n_*_−1_, *R_n_*_−1_, and *S_n_*, respectively. Dashed lines indicate the temporal center of transversal sections shown in c. **c)** Decoding of previous stimulus during previous-trial delay (left), response (middle), and current-trial fixation period (right), for decoders trained in previous-trial delay (mid-delay, 0.5s - 1.0s after *S_n_*_−1_, lower dashed line in b and during previous-trial response (in a window of 0.5s centered on *R*_n-1_, upper dashed line in b. The delay code is stable during delay, disappears during the response, and reappears in current-trial fixation (black line with 95% C.I. error-shading), see also panel d. In contrast, previous-trial information during the response is dynamic and stays around zero in fixation. **d)** Centered average reconstruction of previous stimulus at different epochs for the delay decoder, marked in c (bootstrap-test preferred vs. anti-preferred location: p<0.1e-6, c.i.=[-0.83,-0.27], Cohen’s d=-3.41 (red), p=0.96, c.i.=[-0.29,0.49], Cohen’s d=-0.01 (blue), p=0.002, c.i.=[-0.92,0.13], Cohen’s d=-0.81 (orange), n=15 subjects, shading shows ±s.e.m.). Upper black bars mark significant deviation from zero (bootstrap, p<0.05 in a, p<0.005 in c.

### Reactivation of previous memory information in monkey dlPFC prior to new stimulus presentation

We collected single-unit responses from the dorsolateral PFC (dlPFC) of two monkeys while they performed the task. A substantial fraction of neurons in this area showed tuned persistent delay activity during the mnemonic delay period ^6, 31–34^ (n=206/822). These specific neurons are part of bump attractor dynamics characterizing the memory periods of this task ^6^. Based on our hypothesis that an interplay of activity-silent and attractor mechanisms would support serial biases, we focused our analyses on these neurons, and we grouped them in simultaneously recorded ensembles for decoding analyses (n=94 ensembles, size range 1-6 neurons, Supplementary Fig. 1a).

DlPFC single neuron average firing rates exhibited strong dynamics in the ITI, compared to the stability during mnemonic delay periods (Fig. 1b). Phasic rate increases at response execution (Fig. 1b, *R_n-1_*) and fixation onset (Fig. 1b, *F_n_*) were hallmarks in these dynamics, but we also noticed an increase of firing rate prior to stimulus presentation (Fig. 1b, *S_n_*), which could reflect anticipation of the upcoming stimulus due to fixed length fixation periods. We wondered if these rate changes were also related to dynamical changes in stimulus selectivity. Under the attractor-based hypothesis for serial biases ^28^, sustained stimulus selectivity would be expected to extend from the previous trial’s delay period into the next trial’s fixation period. We measured selectivity by training a linear decoder on spike counts of our small neuronal ensembles, and referenced its accuracy to that obtained by chance using a resampling approach (Methods). During the delay period, neuronal ensembles carried stimulus information and single neurons showed stimulus tuning (Fig. 1c, d red). After report, the memorized location was still decodable from ensemble activity but single neurons’ tuning curves showed a selective suppression of responses in their mnemonic preferred locations (Fig. 1c, green). This could reflect neuronal adaptation mechanisms or else saccade preparation towards the opposite direction to regain fixation. In the middle of the ITI, decoding accuracy was not different from chance and neurons were no longer tuned to the previous stimulus (Fig. 1c,d blue), suggesting that the encoding of the previous stimulus had disappeared from neural activity. However, aligned with anticipatory ramping activity at the end of the fixation period (Fig. 1b), previous stimulus was again decoded just prior to the new stimulus presentation, and single-neuron tuning reappeared (Fig. 1c,d orange). Although the existence of spiking selectivity during the ITI has already been previously reported ^28^, we show here that there is a period in which stimulus information cannot be decoded and then it reappears at the end of the fixation period (*late fixation*). Further, this code in late fixation is a reactivation of the representation active in the previous trial delay. This is supported by 2 pieces of evidence. On the one hand, information reappearance occurred more strongly in those neuronal ensembles that maintained more stimulus information during the delay period (Fig. 1c and Supplementary Fig. 1). Secondly, the converging pattern of noise correlations at the end of the delay ^6^ and in late fixation suggested a similar attractor-like network activation in both periods. Indeed, when the preceding stimulus appeared between two neurons’ preferred locations, these PFC neuron pairs exhibited negative noise correlations in late fixation (Supplementary Fig. 2) – a signature of a fixed-shape bump that diffuses from the initial stimulus location, moving closer to one neuron’s preferred location and away from the other, thus increasing one and decreasing the other neuron’s firing rate^6^. Negative correlations appeared exclusively during late fixation, strongly suggesting a bump reactivation (Supplementary Fig. 2). Taken together, these results support a faithful reactivation of the memory-period representation in the fixation period (*reactivation period*), following a period of absent selective neuronal firing in dlPFC. This reactivation suggests a relationship between mechanisms of delay memory encoding, and mechanisms bridging the ITI to facilitate reactivation prior to the new stimulus.

### Previous trial memory information is reactivated in the fixation period of human EEG

In line with monkey electrophysiology, we found similar previous-trial traces in human EEG (n=15). We extracted alpha power from all electrodes and used a linear decoder to reconstruct the target location from EEG in each trial ^35^ (Methods). The target representation was significantly sustained during delay, response, and the next trial’s fixation period (Fig. 2b, diagonal axis). Considering that EEG is a global measure, this code could be sustained by different representational components (e.g. stimulus, memory, response). We thus trained different linear decoders during delay (500-1000 ms after stimulus onset, *delay code*) and around the response (250 ms before to 250 ms after response, *response code*), and used the respective weights to extract previous-stimulus information throughout different periods of the trial (Fig. 2c). The delay code was stable during stimulus presentation and delay, but disappeared during the ITI, around the time of response. In contrast, the response code did not generalize beyond the time at which the decoder was trained (Fig. 2c). We found that the previous trial’s delay code re-appeared during the fixation period (Fig. 2c, and Fig. 2d, orange), similar to monkey neurophysiology (Fig. 1c). These results provide a confirmatory correspondence with the time-course of mnemonic decoding in the monkey data, but they also show the temporal continuity between qualitatively distinct memory and response codes. The bidirectional transfer of information between memory and response representations in different brain areas could provide a bridge between the memory and ‘reactivation’ periods observed in PFC. Alternatively, response codes may just reflect the output motor commands and mnemonic codes may subsist subthreshold in PFC to allow reactivations. We tested this hypothesis with cross-correlation analysis of PFC units.

### Increased cross-correlation suggests a latent trace during ITI

We sought experimental validation that activity-silent mechanisms in dlPFC still maintained stimulus information during the ITI. We reasoned that if such latent activation (e.g. a synaptic trace ^11^) affected a group of interconnected neurons, these would be more likely to exceed their spiking threshold in synchrony ^10^. Specifically following a preferred cue, neurons would increase their activity in the delay and maintain a latent activity-silent activation in the subsequent ITI that would be reflected in enhanced synchrony, but not enhanced rates. Moreover, we deduced that this reasoning was pertinent only to excitatory interactions: neurons interacting through effective inhibition should instead show reduced spike synchrony following preferred stimuli in the previous trial.

To test this hypothesis, we selected collinear pairs of neurons (distance between the neurons’ preferred locations smaller than 30°, n=67 pairs, Methods) so they had consistent activation (high/low firing rate) in the delay. Following previous studies, we divided the selected pairs based on their whole-trial cross-correlation peak sign in excitatory (*exc*) and inhibitory (*inh*) interactions (refs. ^36, 37^, Methods). We considered two conditions (Fig. 3a, Methods): trials in which the previous stimulus was shown close to either preferred location (*pref*, maximum distance of 40°) or far from preferred locations (*anti-pref*, all other trials). Then, we computed a cross-correlation selectivity index (CCSI) by subtracting the amplitude of the central peak of the jitter-corrected cross-correlation function (coincident spikes within 20 ms, Methods, similar to ref. ^38^) for *pref* and *anti-pref* trials for each neuron pair (Fig. 3b). Our hypothesis predicts positive (negative) CCSI for *exc* (*inh*) pairs in the ITI, i.e. higher (lower) spike synchrony following preferred stimuli. The CCSI computed in a period of the ITI where firing rate had ceased to represent the stimulus (*activity-silent period*, Fig. 1c,d, blue) was positive, reflecting selectivity in neuronal synchrony to the previous stimulus for all interactions (Fig. 3c). We then investigated changes in CCSI for *exc*/*inh* interactions across our two periods of interest: the activity-silent and reactivation periods (Fig 1c, blue and orange, respectively). We found that their reactivation-period CCSI differed significantly, being negative for *inh* interactions and positive for *exc* interactions (Fig. 3c). Finally, we explored the dynamics of selectivity throughout the trial (Fig. 3d): with the exception of immediately after the previous response, where neurons showed anti-tuning to previous-trial stimulus (Fig. 1c), CCSI for *exc* pairs was always positive, indicating stronger central-peak cross-correlation when the previous stimulus was preferred (Fig. 3d, orange). On the other hand, for *inh* interactions CCSI was negative (stronger inhibitory interactions following a preferred stimulus) only during reactivation and the previous-trial delay period (Fig. 3d, green), the periods where PFC firing rates showed stimulus selectivity. This pattern is consistent with the latent memory mechanism residing in excitatory neurons and only being reflected in inhibitory interactions through the collective engagement in bump attractor dynamics, during the delay and at the time of reactivation.

**Figure 3.**
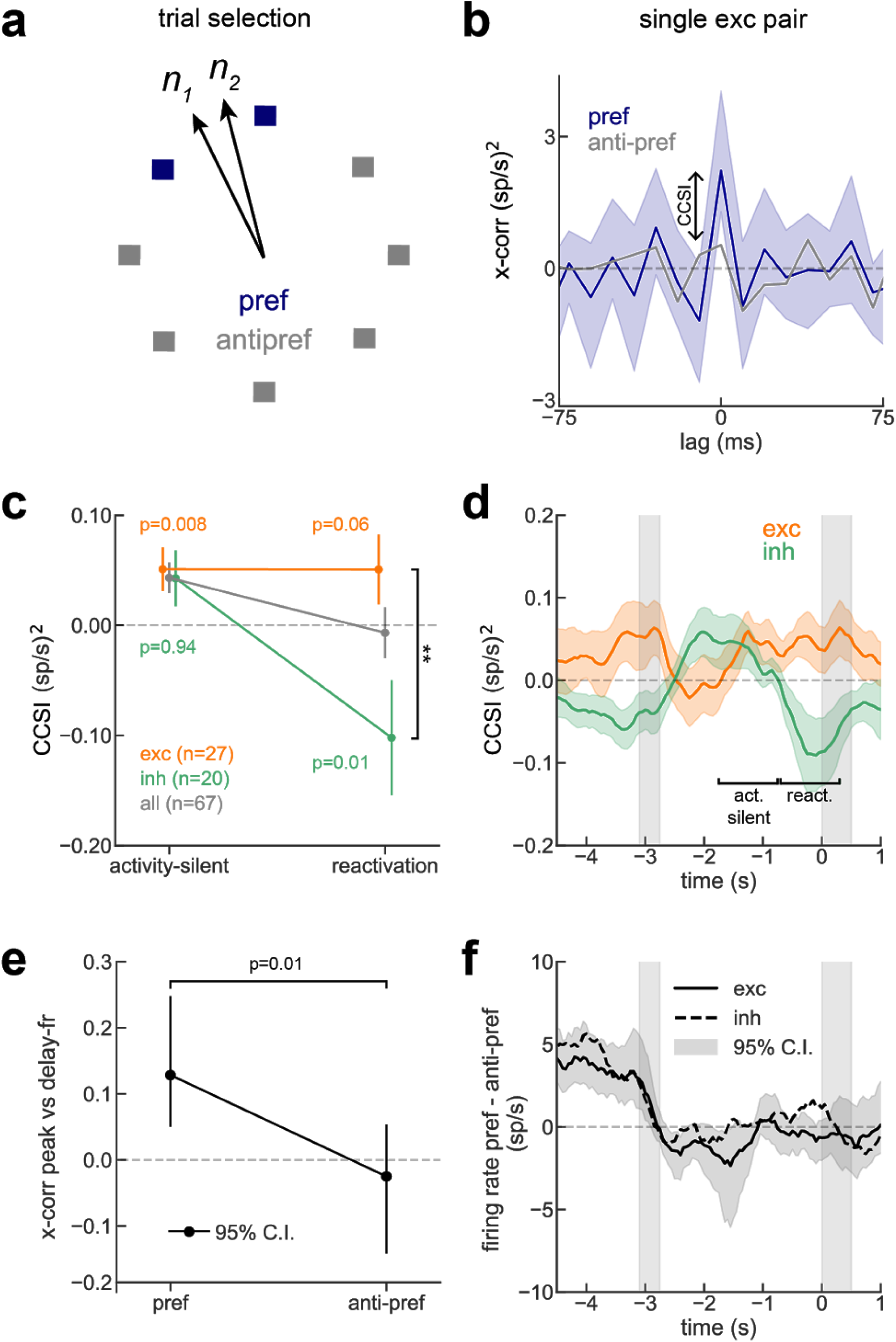
Cross-correlation selectivity to previous-trial stimulus suggests an activity-silent trace in PFC. **a)** Scheme of trial selection: For neuron pairs with similar preferred location (< 60°) we separated trials with stimulus near the pair’s preferred locations (blue, pref) from those where they were far (gray, anti-pref). **b)** Cross-correlation of sample PFC pair shows zero-lag peak selectivity to previous-trial stimulus in activity-silent period (permutation test, p=0.025, Cohen’s d=0.10). **c)** CCSI was consistently positive in activity-silent period, but it became negative for inh interaction pairs in the reactivation period (permutation test, interaction period x exc/inh, p=0.03, Cohen’s d=-0.6). At reactivation, CCSI for *exc* and *inh* differed significantly (**, permutation test, p=0.006, d=0.75). P-values indicated in figure panel report results of one-tailed permutation tests according to our hypotheses (CCSI>0 for *exc*, CCSI<0 for *inh*). **d)** CCSI in the ITI (1-s windows, 50-ms steps) for *exc* and *inh* pairs. Except immediately after report, where neurons show anti-tuning (Fig. 1d), CCSI was positive for *exc* interactions. CCSI was negative for *inh* interactions during previous delay and reactivation. Smoothed with 5-sample square filter. **e)** Trial-by-trial correlation between *exc* pairs’ previous delay spike counts and ITI cross-correlation central peak (activity-silent period in d) is positive only for the *pref* condition (permutation test p=0.017, interaction p=0.01). **f)** Absence of mean firing rate difference between *pref* and *anti-pref* conditions (n=67 pairs) discards confound between rate selectivity and CCSI. Error bars in b and e, 95% C.I; in c and d, s.e.m.

This proves the existence of a latent trace of the stimulus in PFC during the ITI, but it could still be reflecting inputs from a different area. To strengthen the idea that stimulus information is directly transferred from an activity-based to an activity-silent code in PFC, we tested if the selectivity of *exc* interactions during the activity-silent period depended on spiking activity of corresponding neurons in the previous delay period. Assuming a neuron-specific activity-dependent mechanism supporting the activity-silent code in the ITI, we predicted that the magnitude of the cross-correlation central peak in the activity-silent period would correlate on a trial-by-trial basis with the mean spike count recorded in the preceding delay period and specifically for *pref* (and not for *anti-pref*) trials (Methods). This prediction was confirmed in the experimental data (Fig. 3e). Thus, this cross-correlation analysis supports the hypothesis that previous, currently irrelevant stimulus information remains in prefrontal circuits in latent states, undetected by linear decoders that do not take spike timing into consideration (Figs. 1c, 3f).

### Bump-reactivation as a mechanism for stimulus information reappearance

Based on our electrophysiology results and on prior modeling studies ^11^, we formulated the bump-reactivation hypothesis to explain our data. We hypothesized that information held in memory as an activity bump during the previous trial’s delay period ^6^ would be imprinted in neuronal synapses as a latent, activity-silent trace during the ITI. This latent bump could be reactivated by the unspecific anticipatory signal seen in mean firing activity in PFC (Fig. 1b), or anticipatory mechanisms following an external cue that predicts stimulus presentation, such as the onset of a fixation dot (Fig. 2c). In fact, in a separate EEG experiment where fixation lengths were jittered so as to make stimulus onsets unpredictable, we could not find any delay code reactivation (Supplementary Fig. 3).

To test the bump-reactivation hypothesis, we built a bump attractor network model of spiking excitatory and inhibitory neurons. Based on our electrophysiology findings, short-term plasticity (STP) dynamics were included only in excitatory synapses (Methods). In each trial, stimulus information was maintained in activity bumps during the delay, supported by recurrent connectivity between neurons selective to the corresponding stimulus. During the ITI period, model neurons had no detectable tuning to the previous-trial stimulus (Fig. 4a, black line and Fig. 4b, blue) ^22, 27^. However, the synapses of neurons that had participated in memory maintenance in the previous delay were facilitated due to STP (Fig. 4a, blue line). Parallel to our analysis in Fig. 3, this was reflected in the central peak of the ITI cross-correlation for pairs of excitatory model neurons, which maintained selectivity to the previous stimulus (Fig. 4a) even in the absence of single neuron firing rate selectivity (Fig. 4a, blue). We found that single neuron tuning could be recovered from the hidden synaptic trace using a nonspecific input (*drive)* to the whole population (Fig. 4a,c, Methods). Our biologically constrained computational model was thus an explicit implementation of the bump-reactivation hypothesis that we had formulated.

**Figure 4.**
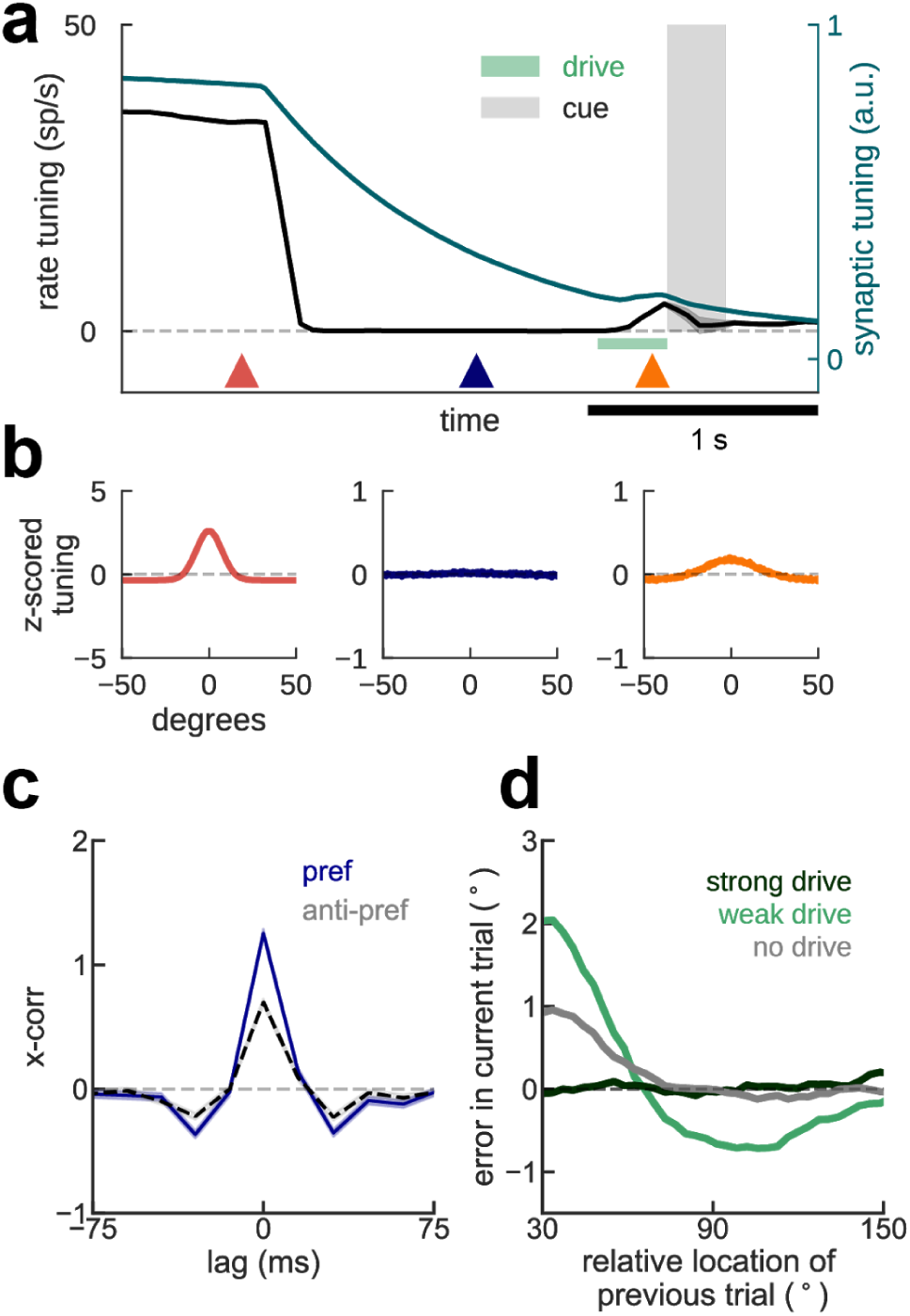
Bump-attractor model with STP accounts for serial dependence and neurophysiology. **a)** Average firing rate tuning (black) and synaptic tuning (green) for 5,000 simulations of two successive trials during delay (Methods). In the mnemonic period (red triangle), both rate and synaptic tuning are at their maximum, both driven by persistent bump-attractor activity (red plot in b). Following the memory period, a nonspecific inhibitory input resets the baseline network state for the duration of the ITI (blue triangle and plot in b). This is reflected in vanishing rate tuning, but long-lasting synaptic tuning that can regenerate firing rate tuning (orange plot in b) through reactivation by a non-specific input drive (light green bar). **b)** Averaged single-neuron tuning to previous-trial stimulus at different epochs marked with colored triangles in a. **c)** Cross-correlation of model neurons in the ITI differed for previous-trial stimulus in the preferred location (pref, blue) and for anti-pref trials (gray) despite no firing rate selectivity (a,b, blue). **d)** Serial bias plots computed from “behavioral responses” (Methods) in 3 different conditions of non-specific excitatory drive. A weak anticipatory drive increases attractive serial biases and produces a repulsive lobe, while a strong drive removes serial biases.

### The impact of bump reactivation on serial biases

We next used our computational model to derive behavioral and physiological predictions to test in our data, in particular in relation to serial biases. In order to simulate serial biases with our computational model, we ran pairs of consecutive trials with varying distance between the two stimuli presented in each simulation. We used the final location of the bump in the second trial (current-trial memory) as the “behavioral” output of the model in that trial. We were able to model the profile of serial biases observed experimentally (Fig. 4d, black; Supplementary Fig. 4), similar to previous models ^22, 27^. To test the impact of bump-reactivation in serial biases, we compared the behavioral output of simulations with and without drive before the second trial stimulus (Methods). Bump reactivation resulted in stronger attractive biases for similar successive stimuli, and in repulsive biases for more dissimilar successive stimuli (Fig. 4d, green line). We found that tuned intracortical inhibition was necessary for this emergence of repulsive biases upon bump reactivation (Supplementary Fig. 4), showing that repulsive biases are caused by repulsive interactions between simultaneously active bumps in the network ^39, 40^, and are absent when there is no reignited bump that recruits localized inhibition at the flanks of the pre-cue bump of activity. We finally tested the dependence of this behavioral effect on the strength of the nonspecific drive. A very short but strong impulse to the whole network during the ITI quickly saturated all the synaptic facilitation variables, effectively removing all serial biases in the output of the network (Fig. 4d, blue). Thus, in this model bump reactivation affects serial biases non-linearly as reactivation strength is varied. In sum, our model can reproduce behavioral and neurophysiological findings described in Figs. 1-3 and derives predictions concerning memory reactivations from silent traces that we then tested in the data.

### Previous stimulus reactivation increases serial biases

The model predicts that higher reactivation of previous memories in the fixation period should be associated with stronger increases in serial biases (Fig. 4d). We tested this prediction in our neural recordings from monkey PFC and EEG recordings on the human scalp.

#### Monkey PFC

We first classified each trial based on leave-one-out decoding of previous stimulus in two different time windows during fixation: during a period with no stimulus information (*activity-silent* period, Fig. 1, blue), and at the time of reactivation (Fig. 1, orange). For each of these 2 windows we separated high-decoding trials (first quartile) from low-decoding trials (all other trials) and computed bias curves separately. We found that serial biases were indistinguishable in the activity-silent period (Fig. 5a) but they were stronger for high-decoding than for low-decoding trials at the time of bump reactivation (Fig. 5b). This follows the prediction of our computational model, assigning behavioral relevance to the bump reactivation prior to stimulus onset. This result was not dependent on a singular selection of trial separations: for different proportions of high- and low-decoding trials the serial bias strengths (Methods) changed smoothly and remained consistent with the reported result (Supplementary Fig. 5). We then repeated the same analysis at different time points of the ITI. A significant difference in serial bias strength (Methods) emerged only when trials were classified as low- vs. high-decoding in the reactivation period (Fig. 1c, 5c, orange), and serial biases remained virtually indistinguishable at all other time points (Fig. 5c).

**Figure 5.**
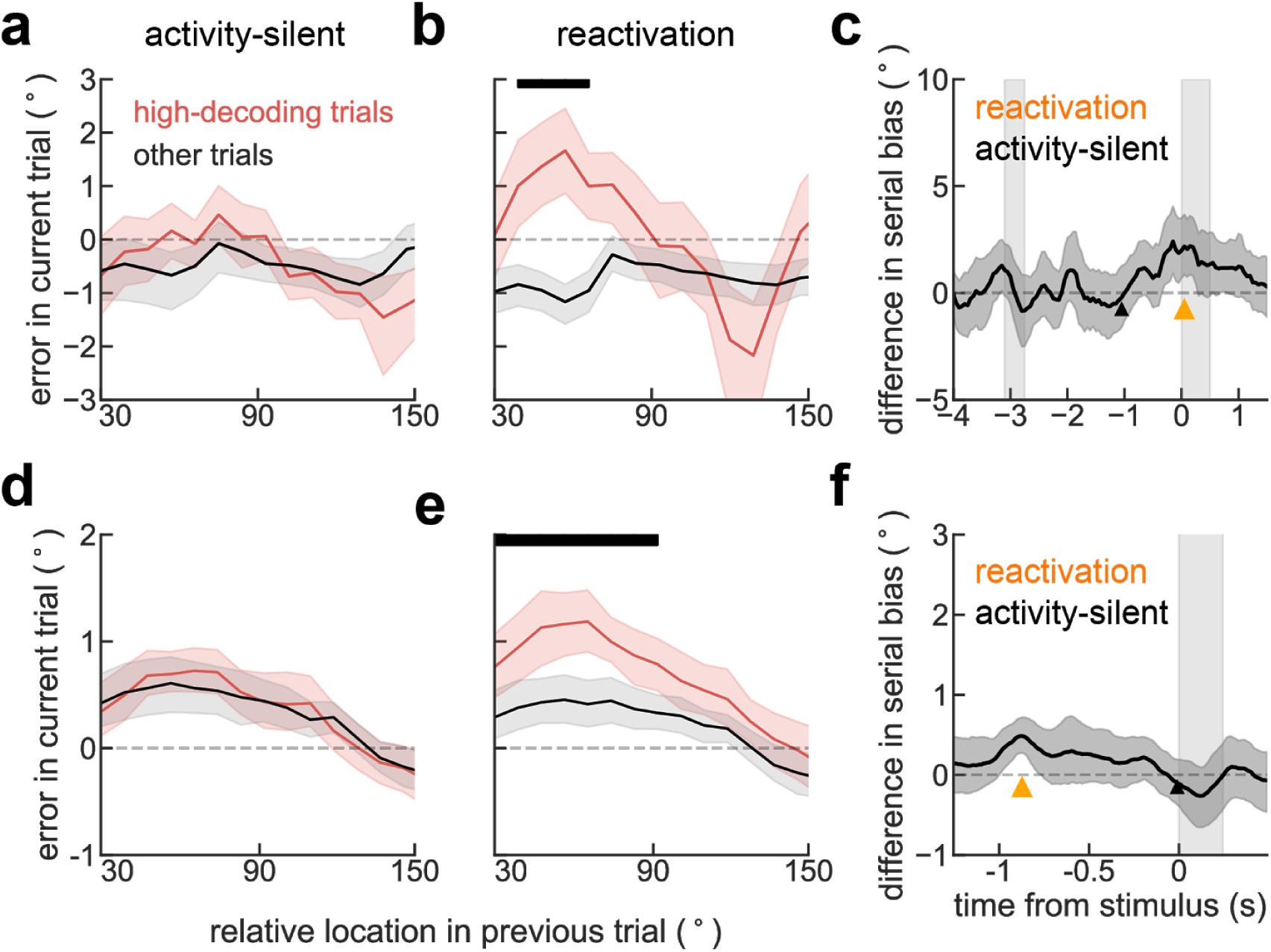
Bump reactivation from a hidden trace increases serial biases. Serial bias curves for trials where previous-trial stimulus information in neurophysiological recordings was high (upper quartile, red) and for all other trials (black), in monkeys (a-c) and in humans (d-f). **a)** Trials selected based on a decoder trained and tested early in the fixation period (black triangle in c), did not reveal differences in serial bias. **b)** Serial biases were markedly enhanced for high-decoding trials when training and testing the decoder at the time of reactivation (Fig. 1c, orange triangle in c). **c)** Difference in serial bias curves between *high-decoding* and *other* trials became significant only at pre-cue, concomitant with reactivation (Fig. 1c). Triangles mark center of 1 s decoding windows for the splits shown in a, b. **d-f)** same analyses for human EEG (n=15). Note that for humans, d corresponds to an activity-silent period in late fixation (black triangle in f), and e to the reactivation period in early fixation (Fig. 2c, orange triangle in f). **f)** As for monkeys, serial bias differences in humans were significant only during reactivation. In c and f, time courses of differences between *high-decoding* and *other* trials were smoothed in time using a 5-sample square filter. Black bars mark significant differences between *high-decoding* and *other* trials (p<0.05, permutation test). Error-bars in c and f, 95% C.I; in a,b,d, e ±s.e.m.

#### Human EEG

Analogous to the analysis performed in monkey data, we grouped trials by their leave-one-out decoding accuracy of the previous stimulus (Methods). We separated high- and low-decoding trials on two different time points: at the time of reactivation (Fig. 2, 5f, orange) and in an arbitrary time point without stimulus information (*activity-silent*, Fig. 5c, black). Consistent with monkey data and our model’s prediction, we found stronger serial bias for high-decoding than for low-decoding trials for the reactivation period (Fig. 5d), but not for the activity-silent period (Fig. 5e), where previous memory content was not decodable (Fig. 2c). The analysis was repeated for all other time points during the fixation period (Fig. 5f). Indeed, behavior exclusively depended on decoding accuracy at the time of delay code reactivation (Fig. 2, orange). Taken together, these results support the hypothesis that previous trial memory reactivation prior to stimulus onset controls serial biases.

### TMS-induced reactivations modulate serial biases

As a causal validation of pre-stimulus PFC reactivation as a mechanism controlling serial biases, we designed a transcranial magnetic stimulation (TMS) perturbation study. This is a relevant experiment because memory-dependent changes in human EEG alpha-power cannot be unequivocally ascribed to a specific brain region, which limits the correspondence of our EEG and monkey dlPFC data. In particular, representations in larger and more organized occipital cortices might contribute strongly to visual EEG signals ^35, 41, 42^, but could yet be driven by top-down projections from association cortices ^43^. Inspired by a previous study that reported reactivation of latent memories using TMS ^16^, we causally tested the implication of dlPFC in serial biases by applying single-pulse TMS during the fixation period. We had two control conditions to test our hypotheses: (1) we targeted the TMS coil at dlPFC and vertex in interleaved blocks; and (2) we randomly chose TMS intensity relative to the subject’s resting motor threshold (RMT) in each trial (*sham:* 0%*, weak-tms:* 70%, and *strong-tms*: 130% of RMT, Methods). We found that TMS modulated serial biases when targeted at dlPFC but not at vertex (Fig. 6). Moreover, our computational model predicted a non-linear dependence with stimulation strength (Fig. 4d), which was supported by the data (Fig. 6b). Interestingly, the behavioral impact of PFC TMS stimulation declined throughout the session, as if subjects desensitized to the TMS pulse (Supplementary Fig. 6). Importantly, we show combined results from two separate experiments of n=10 subjects each, one being a pre-registered replication (Methods, Supplementary Figs. 7, 8). These results provide causal evidence for the involvement of PFC in the serial bias machinery during the ITI. Further, we show that TMS impacts serial biases nonlinearly, as predicted by model simulations that implement the bump reactivation hypothesis via the interplay of bump attractor and activity-silent mechanisms.

**Figure 6.**
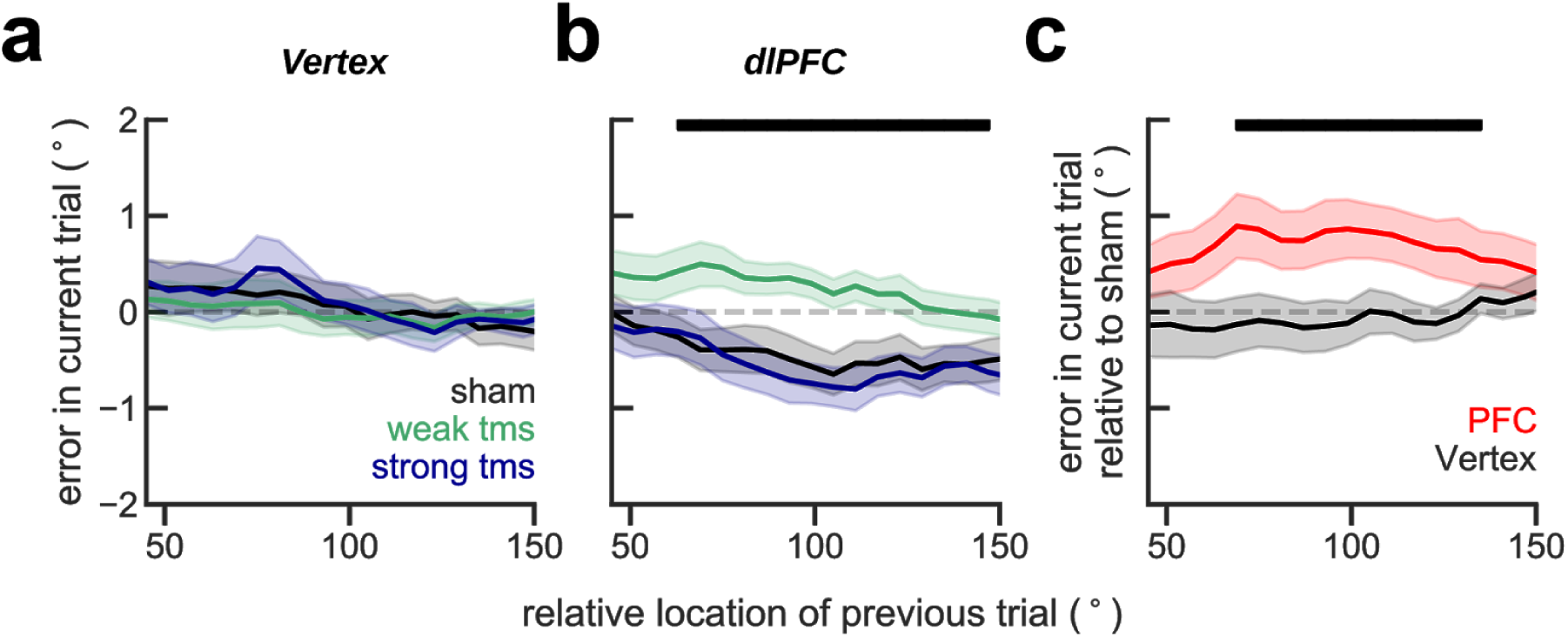
Single-pulse TMS on dlPFC during fixation modulates serial biases non-linearly. Serial bias plots computed in vertex **(a)** and PFC **(b)** blocks, separately for trials with strong fixation-applied TMS pulse (130% of resting motor threshold, RMT, blue), weak (70% RMT, green) and sham (0% RMT, black) for the first 225 trials in each session (n=20 participants, 2 sessions/participant, Supplementary Figs. 6,7,8). Serial biases were modulated by TMS in PFC, but not in Vertex (previous-current stimulus distance (*prev-curr*) × *TMS intensity* × *coil location*, t_18272_=2.21, p=0.027. For dlPFC: *prev-curr* × *TMS intensity*, t_11087_=2.13, p=0.032. For Vertex: t_7166_=0.03, p=0.97. Methods, *Linear mixed models;* analysis performed on the whole session). In PFC, serial bias modulation depended nonlinearly with stimulation strength (ΔAIC=4.6, relative likelihood 0.9, for the comparison of regression models with non-linear vs. linear TMS intensity factor; Methods). **c)** Difference between serial biases computed for sham and weak-tms trials in vertex (black) and in PFC (red) blocks. Error bars are bootstrapped ±s.e.m.. Solid black bars mark significant differences (permutation test, p<0.05).

## Discussion

By studying the neural basis of serial biases, we have shown how the interplay of bump-attractor spiking dynamics and silent mechanisms in prefrontal cortex maintains information in spatial working memory tasks. In these delayed-response tasks, prefrontal tuned persistent activity consistent with bump attractor dynamics characterizes the delay period and correlates with behavioral precision ^6, 44^. We have now seen that this sustained activation disappears from the prefrontal network between trials, but it is reactivated before the new trial (Figs. 1,2) and enhances behavioral serial biases (Figs. 5,6). This reactivation is directly linked to activity recorded in the previous trial: it emerges specifically in those neural ensembles that showed strongest persistent tuning in the delay (Fig. 1c, Supplementary Fig. 1), it is decoded from the human EEG with identical decoders (Fig. 2), and it has the specific fingerprints of bump attractors as evaluated with pairwise spike-count correlations (Supplementary Fig. 2). Activity-silent mechanisms in prefrontal cortex bridge these two disconnected periods of persistent activity, carrying stimulus selectivity from one trial to the next (Fig. 3). Importantly, this latent tuning is directly associated with trial-by-trial spiking activity in the preceding delay period (Fig. 3e), thus establishing a coupling between activity-based and activity-silent mechanisms in PFC. Taken together, our results are consistent with the view that attractor-based and activity-silent mechanisms are jointly represented in the local prefrontal circuit and that their tight interplay supports behavior in spatial working memory. We specified this in a computational network model: delay-period attractor dynamics imprint activity-silent mechanisms, which then retain information between trials and allow reactivations to recapitulate attractor states (Fig. 4).

Our data provides experimental support that non-specific PFC stimulation can revive subthreshold information, similar to the modeling ideas put forward in ref. ^11^. While a recent study ^28^ found neural firing selectivity from previous cues bridging brief ITIs in the frontal eye field, our study, involving longer ITIs, did not find such continuous selectivity in PFC single-neuron firing rates. Rather, information was still present in synchrony parameters, thus revealing a latent subthreshold tuning, and this tuning could be reinstated in firing rates when triggered by external events, as previous neuroimaging and EEG studies have speculated ^16, 45–47^. However, our data also supports the idea that activity-silent and attractor-based mechanisms are not orthogonal, alternative mechanisms but they are largely co-expressed in the circuit and underlie different behaviors: whereas persistent attractor-based activity is engaged during active maintenance of memories, activity-silent maintenance supports secondary, possibly involuntary memory traces, here expressed in small serial biases. Similar ideas have been proposed in the context of attended and unattended memories ^16, 29, 30, 45^. Note, however, that in our proposed framework the close interplay between attractor-based and activity-silent mechanisms does not allow activity-silent memories to be protected from intervening attractor-based activations in the circuit. This yields the prediction that serial-bias-like patterns of interference between unattended and attended memories should be observed in these experiments.

Critically, we demonstrated that the proposed mechanisms can be directly linked with behavior: we found robust evidence for the role of bump reactivations from activity-silent traces in generating working memory serial biases (Fig. 5). This explicit demonstration of the possible behavioral impact of activity-silent traces is an advance over previous studies ^36, 48^. Note however that recent causal evidence in rodents ^49^ may also be interpreted as revealing the impact of parietal reactivations in history-dependent biases. Direct evidence could be sought with activation optogenetics at different time points in the ITI, which should increase biases. In addition, we obtained explicit causal evidence supporting the role of reactivations prior to the trial in enhancing serial biases by using a TMS perturbation approach and validating the predicted behavioral effects (note the pre-registered replication, underscoring the robustness of the results, Methods, Supplementary Figs. 6-8). Previous single-pulse TMS studies in working memory have failed to find strong behavioral effects when pulses were applied in the delay period ^16^. Our application of pulses in the possibly quiescent fixation period may have enhanced the effect of the TMS pulse and facilitated its impact on behavioral reports. Our TMS experiment also clarified our EEG results by demonstrating the role of prefrontal cortex in serial biases. Because we did not concurrently acquire EEG during the TMS study, we could not directly measure the stimulus representation reactivation induced by the TMS pulse. However, prior work has shown the reactivation of EEG memory representations with TMS^16^, albeit in different conditions (pulses in the memory period). Intriguingly, serial biases for trials without TMS stimulation in PFC-stimulation blocks were repulsive (Figure 6b). We speculate that this was due to suppressive carry-over effects in PFC from previous TMS-stimulated trials in the block. In fact, studies combining TMS with single unit recordings report that a fast, excitatory TMS effect is often followed by a long-lasting inhibitory effect ^50, 51^. Future work involving more fine-grained TMS intensities and carefully controlled block designs will be necessary to clarify these results further.

We proposed a computational model that can parsimoniously explain our data using short-term facilitation in the synapses of a recurrent network. Short-term plasticity has also been used in previous computational models of interacting activity-based and activity-silent dynamics ^11, 12, 15^ and of serial biases ^22, 27^. Beyond previous modeling efforts, we explored the mechanistic requirements of code reactivations prior to a new trial, and we derived predictions whose validation conferred plausibility to the model. Of course, our findings do not unequivocally identify this mechanism and we could have chosen another mechanism with a long time constant to implement our hypothesis computationally (e.g. calcium-activated depolarizing currents ^52^, depolarization-induced suppression of inhibition ^13^, short-term potentiation ^53^, etc). Still, several lines of evidence support the involvement of short-term plasticity in prefrontal function. First, there is explicit evidence for enhanced short-term facilitation and augmentation among PFC neurons in *in vitro* studies ^54, 55^. Second, extracellular recordings in behaving animals cannot directly probe activity-silent mechanisms, but indirect evidence for synaptic plasticity has been gathered in cortical activity correlations of animals engaged in working memory tasks ^36, 48^. Our study also follows this approach to seek evidence for activity-silent stimulus encoding, but we apply it specifically at time periods without firing rate codes for task stimuli, thus unambiguously decoupling activity-silent from activity-based selectivity.

In sum, we provide experimental evidence that subthreshold traces of recent memories remain imprinted in PFC circuits, and influence behavioral output in working memory in particular through network reactivations of recent experiences. Our findings suggest that the dynamic interplay between attractor and subthreshold network dynamics in PFC supports closely associated memory storage processes: from effortful memory to occasional reactivation of fading experiences.

## Acknowledgments

This work was funded by the Spanish Ministry of Science, Innovation and Universities and European Regional Development Fund (Refs: BFU2015-65315-R and RTI2018-094190-B-I00); by the Institute Carlos III, Spain (grant PIE 16/00014); by the Cellex Foundation; by the Generalitat de Catalunya (AGAUR 2014SGR1265, 2017SGR01565); and by the CERCA Programme/Generalitat de Catalunya. CC was supported by NIH grant R01 EY017077. JB was supported by the Spanish Ministry of Economy and Competitiveness (FPI program). HS was supported by the “la Caixa” Banking Foundation (Ref: LCF/BQ/IN17/11620008), and the European Union’s Horizon 2020 Marie Skłodowska-Curie grant (Ref: 713673). KA was supported by NIH grant T32-MH020002. This work was developed at the building *Centro Esther Koplowitz*, Barcelona. We thank D Lozano-Soldevilla for assistance with EEG analyses, LCG Del Molino for valuable insights during the development of early versions of the model, and Alfonso Renart and Jaime de la Rocha for their comments on the manuscript.

## Data availability

All data that support the findings of this study are available from the corresponding author upon request.

## Code availability

The custom code is available from the corresponding author upon request.

## Author contributions

JB and AC performed monkey data analysis. JB and AC developed the model. HS and AC designed human research. HS and AG performed human experiments. HS, JB and AC performed human data analyses. KA performed preliminary human data analyses. JB, RM, JV and AC designed the TMS experiments. RM performed TMS experiments. SL performed monkey experiments. CC designed monkey research. JB, HS and AC discussed the results and wrote the manuscript. All authors revised the manuscript and gave critical comments.

## Competing Interests

The authors declare no competing interests.

## Materials and Methods

### Behavioral task and recordings

#### Monkey behavioral task and recordings

Four adult, male rhesus monkeys (*Macaca mulatta*) were trained in an oculomotor delayed response task requiring them to fixate, view a peripheral visual stimulus on a screen at a distance of 50 cm and make a saccadic eye movement to its location after a delay period. During execution of the task, neurophysiological recordings were obtained from the dorsolateral prefrontal cortex (dlPFC). Detailed methods of the behavioral task, training, surgeries and recordings, as well as descriptions of neuronal responses in the task have been published previously ^6, 31–34^ and are only summarized briefly here. Visual stimuli were 1° squares, flashed for 500 ms at an eccentricity of either 12° or 14°. Stimuli were presented randomly at 1 out of 8 possible locations around the fixation point. A delay period lasting 3 s followed the presentation of the stimulus, at the end of which the fixation point turned off, and a saccade terminating within 5° from the location of the remembered stimulus was reinforced with liquid reward. Although fixation was maintained through cue and delay periods, we denote “fixation period” the interval between fixation onset and cue onset, when the only behavior expected was fixation (*fixation period*, Fig. 1b). A fixed inter-trial interval (ITI) of 3.1 seconds elapsed between fixation cue extinction and the start of a new trial with fixation onset (*ITI*, Fig. 1b). Eye position was monitored with a scleral eye coil system in two monkeys and an ISCAN camera in the other two. From 2 of those monkeys, we collected single-unit responses from dlPFC using tungsten electrodes of 1–4-MΩ impedance at 1 kHz, while they were performing the task. Simultaneous recordings were obtained by arrays of 2-4 microelectrodes, spaced 0.2 – 1 mm apart of each other. A substantial fraction of neurons in this area showed tuned persistent delay activity during the mnemonic delay period of the task (n=206/822, ^6, 31–34)^. For decoding analyses, we grouped those neurons in simultaneously recorded ensembles (total of n=94 neural ensembles, 1-6 neurons per ensemble, Supplementary Fig. 1a). All experiments were conducted in accordance with the guidelines set forth by the US National Institutes of Health, as reviewed and approved by the Yale University Institutional Animal Care and Use Committee.

#### Human participants and behavioral task

Thirty-five (35) neurologically and psychologically healthy volunteers with normal or corrected vision (EEG experiment n=15 (4 male), 21.27 ± 4.86 years, (mean ± std); TMS experiments n=20 (6 male), 29.86 years ± 9.55 years (mean ± std)) from the Barcelona area provided written informed consent and were monetarily compensated for their participation, as reviewed and approved by the Research Ethics Committee of Hospital Clínic (Barcelona). During both EEG and TMS experiments, each participant performed two sessions of approximately 1.5 h. To perform behavioral and EEG analyses, we concatenated the two sessions for each subject. Stimuli were presented on a 17’’ HP ProBook using Psychopy (version 1.82.01) viewed at a distance of 65 cm. The TMS study consisted of a first experiment with 10 subjects, and a pre-registered replication experiment (https://osf.io/rguzn/) with 10 more subjects (Supplementary Figs. 6-8). For all 3 studies (one EEG and two TMS experiments), we recruited independent subject pools.

In each 1.5 h EEG session, participants completed 12 blocks of 48 trials (except for one participant, who completed 12 blocks in one, and 11 blocks in the second session). Each trial began with the presentation of a central black fixation dot (.5 × .5 cm) on a gray background. After 1.1 s of fixation, a single colored circle (stimulus, diameter 1.4 cm) appeared for .25 s at any of 360 circular locations at a fixed radius of 4.5 cm, randomly sampled from a uniform distribution. In 66.67% of trials (a total of 768 trials per subject), the stimulus was followed by a 1 s delay in which only the fixation dot remained visible. In the remaining trials, the delay duration was either 3 s (16.67% of trials, 192 trials per subject) or 0 s (16.67% of trials, 192 trials per subject). Trials with 0 s delay were excluded from the analyses in this study. The change of the fixation dot color (from black to the stimulus color) instructed participants to respond (response probe). Participants responded by making a mouse click at the remembered location. A transparent circle with a white border indicated the stimulus’ radial distance, so the participant was only asked to remember its angular location. After the response was given, the cursor had to be moved back to the fixation dot to self-initiate a new trial (median 1.5 s). Participants were instructed to maintain fixation during pre-stimulus fixation, stimulus presentation, and delay and were free to move their eyes during response and when returning the cursor to the fixation dot. Colors (1 out of 6 colors with equal luminance) were randomly chosen with equal probability for each trial.

Stimuli and trial structure in the TMS task were similar to the EEG task, except for the fixation period duration (0.6 s) screen background (white), stimulus color (black), and response probe color (red). At the end of the fixation period (16.7 ms prior to stimulus onset), a single TMS pulse was applied in half of vertex trials (weak/strong tms or sham trials), and in two thirds of prefrontal trials (weak, strong or sham trials). See TMS details below. Only delays of 1 s were used in this experiment. Participants completed 4 blocks of 90 (vertex) and 4 blocks of 130 (PFC) trials within each session. In the first TMS study, these 8 blocks were randomly shuffled for each session. In the replication TMS study, we successively alternated vertex and PFC blocks within each session, and the 2 sessions of a given participant started alternatively with each area in a counterbalanced design.

#### EEG recordings and preprocessing

We recorded EEG from 43 electrodes attached directly to the scalp. The electrodes were located at Modified Combinatorial Nomenclature sites Fp1, Fpz, Fp2, AF7, AFz, AF8, F7, F3, Fz, F4, F8, FT7, FC3, FCz, FC4, FT8, A1, T7, C5, C3, Cz, C4, C6, T8, A2, TP7, CP3, CPz, CP4, TP8, P7, P3, Pz, P4, P8, PO7, PO3, POz, PO4, PO8, O1, Oz and O2. Sites were referenced to an average of mastoids A1 and A2 and re-referenced offline to an average of all electrodes. We further recorded horizontal EOG from both eyes, vertical EOG from an electrode placed below the left eye and ECG to detect cardiac artifacts. We used a Brainbox® EEG-1166 EEG amplifier with a .017-100 Hz bandpass filter and digitized the signal at 512 Hz using Deltamed Coherence® software (version 5.1).

We visually identified and excluded outlier trials based on each trial’s EEG summary statistics (variance and kurtosis of samples within segmented periods of time). To reduce artifacts in the remaining data, we ran an independent component analysis (ICA) on the trial-segmented data and corrected the signal for blinks, eye movements, and ECG signals, as identified by visual inspection of all components. Data were Hilbert-transformed (using the FieldTrip function “ft_freqanalysis.m”) to extract frequencies in the alpha-band (8-12 Hz) and total power was calculated as the squared complex magnitude of the signal. Finally, we excluded trials in which lognormal alpha-power at any electrode exceeded the time-resolved trial average of lognormal alpha-power by more than 4 standard deviations. In total, we rejected an average of 1.87% ± 1.24% (mean ± std) of trials per participant. Excluding rejected trials and trials with 0 sec delay, we used 935.12 ± 29.03 trials per participant. To concatenate data from the two sessions for the same subject, we normalized each session’s alpha-power for each electrode separately.

#### Transcranial Magnetic Stimulation

Stimulation was performed in the TMS study using a Magstim Rapid 2 machine with a 70 mm figure-of-eight coil. TMS target points were located using a BrainSight navigated brain stimulation system that allowed coordination of the coil position based on the participant’s structural MRI (sMRI) scan. A region of interest in dlPFC was defined using NeuroSynth ^56^ term-based meta-analysis of 53 fMRI studies associated with the key phrase ‘spatial working memory’. This mask was transformed into each subject’s sMRI space. Vertex target points were defined using 10-20 measurement system. Stimulator intensity, coil position, and coil orientation were held constant for each participant for the duration of each session. In order to mask the sound of TMS coil discharge, we had participants listen to white noise through earphones for the duration of the session. White noise volume was selected based on participant threshold for detecting TMS click using the staircase method (2-up, 1-down). Stimulation intensity was determined by the individually-defined resting motor threshold (RMT). We applied 2 different TMS intensities at 70% RMT (weak-tms) and 130% RMT (strong-tms), depending on the trial (see text). To reduce the number of trials per session, we applied strong-tms at Vertex in the original study, but weak-tms for the replication study (pre-registered at https://osf.io/rguzn/, Supplementary Figs. 7 and 8). The stimulation parameters were in accordance with published TMS guidelines ^57^. In a post-experiment debriefing session, we collected information about the subjective experience of the participants. Many participants (13 out of 20) reported facial muscle twitching in dlPFC blocks. This is an unlikely explanation for the effects observed in Fig. 6 because (1) twitching is expected to increase with TMS intensity but we instead observed a non-linear dependency in our effect (Fig. 6b), and (2) behavioral performance in our task as measured by the precision of the responses was not modulated by TMS intensity in dlPFC blocks (linear mixed model as described below: θ_e_ ∼ *intensity* + (1|*subject*), p>0.5), suggesting that our reported intensity-dependent effect (Fig. 6b) was not the result of a general behavioral impairment caused by facial twitching.

### Serial bias analysis

#### Human

For each trial, we measured the response error (θ*_e_*) as the angular distance between the angle of the presented stimulus and the angle of the response. To exclude responses produced by guessing or motor imprecision, we only analyzed responses within an angular distance of 1 radian and a radial distance of 2.25 cm from the stimulus. Further, we excluded trials in which the time of response initiation exceeded 3 s, and trials for which the time between the previous trial’s response probe and the current trial’s stimulus presentation exceeded 5 s. On average, 2.99% ± 4.51% (mean ± std) of trials per participant were rejected.

We measured serial biases as the average error in the current trial as a function of the circular distance between the previous and the current trial’s target location (θ*_d_*) in sliding windows with size π/3 and in steps of π/20. To increase power and correct for global response biases, we calculated a ‘folded’ version of serial biases as follows ^58^. We multiplied trial-wise errors by the sign of θ*_d_* : θ′*_e_* = θ*_e_* _*_ *sign*(θ*_d_*), and used absolute values of θ*_d_*. Positive mean folded errors should be interpreted as attraction towards the previous stimulus and negative mean folded errors as repulsion away from the previous location. For difference in serial bias analyses (Fig 5f), we averaged folded errors for close prev-curr distances (between 0 and π/2).

#### Monkey

In contrast with the human experiments, the stimulus distribution was discrete for all the monkey experiments. On each trial, the subject was cued to 1 of 8 possible cue locations equidistant on a circle. This restricted the minimal angular distance between cues in two consecutive trials to be π/4. To have a finer resolution to calculate serial biases, we capitalized on the response variability on each trial: we computed θ*_d_* as the distance between the current trial’s stimulus and the previous trial’s response (instead of the previous trial’s stimulus). Similar methods to humans were used, except for Fig. 1a, where we used smaller sliding window sizes (π/10 in steps of π/100), essential to capture the thinner attractive serial bias profile in monkeys (Fig. 1a). We attribute specific differences in our monkey and human serial bias curves (Fig. 1a, 2a) to our specific monkey samples, as studies with larger samples (5+2) have reported monkey behavioral biases more consistent with the human literature ^24, 28^.

### Statistical methods

Unless stated otherwise, all hypothesis testing were two-tailed (permutation tests or bootstrap hypothesis test, n=10^6^) and confidence intervals (c.i.) are at [2.5, 97.5] percentiles of a bootstrapped distribution. Using bootstrap distributions, we avoid assuming normality for our statistical tests. One exception was the linear model used for TMS data analyse, in which normality was assumed. Supplementary Fig. 9 shows the distribution of residuals of this model and corresponding qqplot. There was a significant deviation from normality in extreme values. This did not compromise our statistical inference, because of the large sample size (n= 18299) ^59^ and because the interaction of interest was confirmed by model-free analyses (Fig. 6, Supplementary Fig. 6-8).To test the effect of TMS on serial biases, we fit a linear mixed-effects model using the R function *lme* ^60^. In particular, we modeled trial-wise behavioral errors θ*_e_* as a linear model with interaction terms for *coil location (PFC vs. vertex)*, TMS *intensity (strong-tms, sham, and weak-tms)* and the sine of θ*_d_* (*prev-curr*), which approximates the expected dependency of θ_e_ on θ*_d_* in the presence of serial biases (θ*_e_* ∝ *sin*(θ*_d_*)). We incorporated the non-linear dependency of serial bias on stimulation intensity that our model simulations predicted, by using −1, 0 and 1 for strong-tms, sham and weak-tms, respectively. In one model, we used instead the nominal percent of RMT TMS intensity used (70, 0, 130, respectively) for comparison (Fig. 6b). We accounted for subject-by-subject variability by including random-effect intercepts and random-effect coefficients of *prev-curr*. The full, three-way interaction model was:

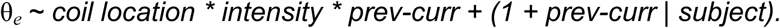

### Decoding stimulus information

#### Monkeys

##### Population decoder

For each recorded ensemble, we decoded stimulus θ*_j_* in trial *j* by modelling it as a linear combination of the spike counts of simultaneously recorded neurons *n_i_*, computed in sliding windows of 0.5 s and steps of 0.1 s during that trial:

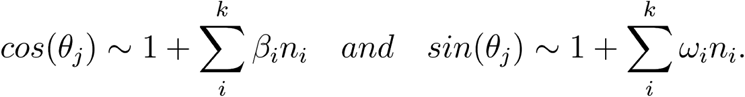

For each set of neurons, we trained two sets of weights β*_i_* and ω*_i_* on 80% of randomly selected trials and tested in the remaining trials. We applied Monte-Carlo cross-validation with 50 random splits to obtain angle estimates 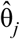. We obtained a measure of error (*err*) by averaging across splits the mean absolute error 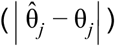 in each split.

##### Accuracy of ensembles: Distance from shuffle

To establish the significance of decoding accuracy (*z*), we compared each ensemble’s decoding error (*err*), to the distribution of decoding errors in 1,000 shuffled stimulus sequences (*err_s_*). By shuffling the list of stimuli presented in the particular recording of each ensemble, we maintained the characteristics of the distribution (e.g. unbalanced distribution of stimuli), but effectively destroyed correlations between stimuli and neural activity.

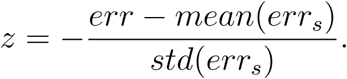

In Fig. 1c and Supplementary Fig. 1b, we tested separately ensembles that had the strongest and weakest decoding accuracy in the delay by obtaining *z* from spike counts in the delay period, and classifying the ensembles based on *z* : ensembles within the top tertile (*high-decoding delay* ensembles), and those in the bottom tertile (*low-decoding delay* ensembles).

##### Accuracy of single trials: Leave-one-out decoder

To measure stimulus information on a trial-by-trial basis, we used leave-one-out cross-validation. We regressed the β*_i_* and ω*_i_* weights in all trials, except the one left out for testing. For these analyses we computed spike counts in windows of 1 s in steps of 50 ms.

#### Humans

##### Linear decoder

EEG alpha power is known to decrease in occipital sites contralateral to attended locations and locations being actively maintained in working memory ^35, 41, 42, 61^. We used this feature to decode the stimulus’ angular position from the distribution of alpha power over all 43 electrodes. We trained the decoder on the previous trial’s stimulus label and decoded this information throughout previous and current trial. Trialwise alpha power for each electrode was modeled as a linear combination of a set of regressors representing the stimulus location in the corresponding trial, *U* = *WM*, where *U* is a *J* × *K* matrix of alpha power measured at electrode *j* in trial *k*, *M* is the *N* × *K* design matrix of values for regressor *n* in trial *k*, and *W* is the *J* × *N* weight matrix, mapping the weight for regressor *n* to electrode *j*. *U* and *M* (determined by the stimulus, see below) were given by the experiment, while *W* was fitted using least squares (see below).

##### Design matrix M

The design matrix *M* is a set of eight regressors *M_n_* representing expected “feature activations” ^62^ for feature *n* in trial *k*. The value of regressor *M_n_* in trial *k* was determined as |*sin*(*n*π/8 − *s_k_* π/8 + π/2) ^7^|, where *s_k_* = [0 .. 7] indicates which one of eight angular location bins (width π/8 rad) included the stimulus shown in trial *k*.

Similar to monkey analyses, we measured single-trial stimulus representations using leave-one-out cross-validation, ensuring equal number of trials from each location bin in the training set (*U_t_* and *M_t_*). We estimated the weight matrix Ŵ and the left-out trial *k*’s design matrix *M_k_* by:

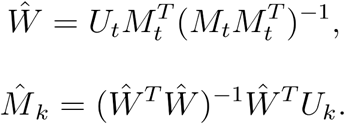

For each trial and time point, we repeated this analysis 100 times with randomly chosen training sets (except for the temporal generalization matrix, for which 10 repetitions were run, Fig 2b), and averaged 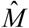 over all repetitions. Finally, we estimated the predicted angle 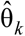 as the direction of the vector sum of feature vectors with length 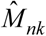 pointing at angular location bin centers *b_n_* = *n*π/8 (*n*=1..8). Trialwise decoding strength was then defined as 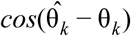.

##### Cross-temporal decoding

To explore the temporal generalization of the mnemonic and the response code over time, we trained decoders in independent time windows of the previous and current trial, and tested them in all time points of consecutive trials (from .25 s to 1.25 s after previous stimulus onset (Fig. 2c, left), -.25 s to .25 s after previous response (Fig. 2c, middle), and −1.25 s to .25 s after the current trial’s stimulus onset (Fig. 2c, right)). For the temporal generalization matrix (Fig. 2b), we averaged training and test data over 50 samples (≈ 97.77 ms). High-resolution time courses of mnemonic and response code (Fig. 2c) were obtained by training the decoder on averaged data from 0.5 s to 1 s after previous stimulus onset and −.25 s to .25 s relative to the response time (dashed lines in Fig. 2b), respectively, and by testing on averaged data from five samples (≈ 9.77 ms) through consecutive trials.

### Preferred location

We computed the preferred locations of each neuron. Similar to ref. ^6^, preferred location was determined by computing the circular mean of the cue angles (0° to 315°, in steps of 45°) weighted by the neuron’s mean spike count over the delay period (3 s) following each cue presentation.

### Cross-correlations

#### Dataset

For the estimation of functional connectivity we estimated cross-correlations by computing the jittered cross-covariances ^63^ of spike counts from simultaneously recorded neuron pairs, whose preferred locations where separated by a maximum of 60° (n=67). We included pairs of neurons recorded from the same electrode (n=21) and pairs recorded from different electrodes (n=46), and we confirmed that the results held when analyzing only pairs from different electrodes (Fig. 3c, *exc* p=0.01, n=20; *inh* p=0.04, n=13, one-sided permutation test). For each pair we selected those trials where the presented cue fell within the preferred range (*pref*, within 40° from either preferred locations) or outside the preferred range (*anti-pref*, all the other trials). We discarded those trials without at least 1 spike for each neuron in the pair.

#### Jittered cross-covariance

We used the Python function scipy.signal.correlate to compute cross-covariances between spike trains of simultaneously recorded pairs. Spikes were counted in independent windows of 10 ms ^38, 64^. For each trial, 1000 jittered cross-covariances were computed as follows ^63^. We shuffled the spike counts within non-overlapping windows of 50ms and computed cross-covariance for each of these jittered spike counts. This captured all the cross-covariance caused by slow dynamics (>50ms) but destroyed any faster dynamics. Finally, we removed the mean of these jittered cross-covariances from each trial’s cross-covariance, ending up with correlations due to faster dynamics (<=50ms). We considered the magnitude of the central peak of the cross-covariance in our analyses by averaging 3 bins (+1/-1 bin from the zero-lag bin). For the time resolved cross-correlation function (Fig. 3c,d), we repeated this process for sliding windows of 1 s and steps of 50ms, and averaged across trials and neuronal pairs.

#### Putative excitatory and inhibitory interaction

Similarly to ref. ^36^, based on the average central peak of the cross-correlation function in the whole trial [-4.5 s, 2.5 s], we classified each pair into 3 subgroups: 1) those with positive peak for both preferred and anti-preferred trials were classified as putative *excitatory* interactions (*exc*), 2) those with negative peak for both preferred and anti-preferred trials were classified as putative *inhibitory* interactions (*inh*) and 3) we discarded those with inconsistent peak sign between *pref* and *anti-pref* trials. In total, we analysed the cross-correlation time course of n=47 pairs of neurons (n=27 *exc* and n=20 *inh*; from the same electrode n=20 *exc* and n=13 *inh*).

#### Delay rate vs ITI cross-correlation analyses

In Fig. 3e we sought evidence for an interplay between attractor and subthreshold network dynamics in PFC. To this end, we computed the trial-by-trial correlation between the cross-covariance peak (see above) in the ITI - at a time point when there was no firing rate tuning (*activity-silent* period, Fig. 3d) - and the mean firing rate of the two neurons at the end of the preceding delay period (last 2 seconds, *delay-fr*, Fig. 3e), for *exc* interaction pairs under the *pref* and *anti-pref* condition (see above). A local activity-dependent subthreshold mechanism for ITI memory traces predicts that for *pref* trials, but not for *anti-pref* trials, firing rate variations in the delay period determines the degree of latent variable loading (cross-covariance peak) in the ITI (Fig. 3e).

### Simulating bump reactivation

We used a previously proposed computational model ^39, 65, 66^ to study serial dependence between two consecutive trials. The model consists of a network of interconnected 2048 excitatory and 512 inhibitory leaky integrate-and-fire neurons ^67^. This network was organized according to a ring structure: excitatory and inhibitory neurons were spatially distributed on a ring so that nearby neurons encoded nearby spatial locations. Excitatory connections were all-to-all and spatially tuned, so that nearby neurons with similar preferred directions had stronger than average connections, while distant neurons had weaker connections. All inhibitory connections were all-to-all and untuned. Network parameters were taken from (Compte et al. 2000) except for:

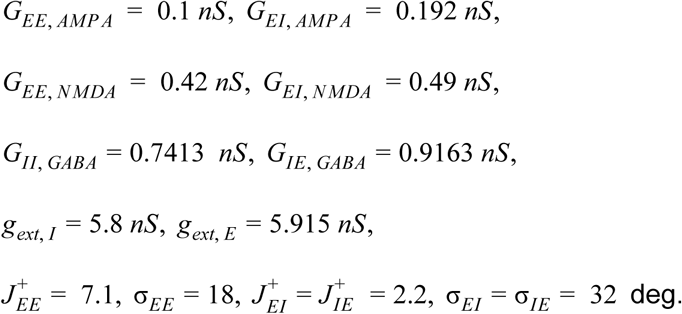

#### Short-term plasticity

Simulation of “activity-silent” mechanisms during the inter-trial period, was done by adding two more variables *x* and *u*, as described in refs. ^11, 68^, to excitatory presynaptic neurons:

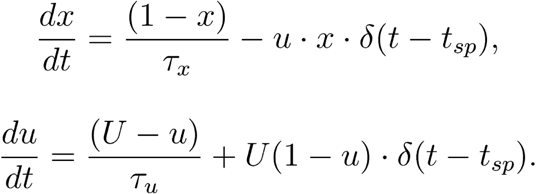

With *t*_sp_ marking all spike times and δ(*t*) being the Dirac delta function. We used parameters *U* = 0.2, τ *_x_* = 0.2 *ms*, τ *_x_* = 1500 *ms*. The effective conductance of each excitatory synapse was then *g*⋅*u*⋅*x*, with *g* being the corresponding maximum conductance parameter (see above). These short-term plasticity dynamics affected only AMPAR-mediated recurrent connections in the network.

#### Stimulation and behavioral readout

External stimuli were fed into the circuit as weak inputs (0.25 nA) to neurons selective to the stimulus as described in Compte et al. (2000). Each simulation of our computational model consisted in two trials run in sequence: a first stimulus of duration 250 ms, a first delay period of 1000 ms, a network resetting input (nonspecific current −2.61e-10 nA, duration 300 ms), an intertrial interval of duration 1300 ms, a second stimulus (250ms) and a second delay period of 1000 ms. The first and second cue stimuli were independently drawn randomly from 360 uniformly distributed angular values, and only the network readout of the second trial was analyzed to obtain a “behavioral readout”. The readout was obtained with a “bump tracking” procedure: starting at cue presentation the instantaneous network readout was derived as the angular direction of the population vector of single-neuron firing rates (computed in windows of 250 ms, sliding by 100 ms) considering the ±100 neurons surrounding the readout estimated in the previous time step. The instantaneous readout was iteratively derived to track the center of the bump (and ignoring possible elevated activity extending from the fixation period) and the final behavioural output was defined as the readout in the last 250 ms of the trial. Serial bias was calculated by measuring single-trial errors (behavioral readout minus target location) in relation to previous-current distance of stimulus cue values, as described above for experimental data.

#### Consecutive trials and re-ignition

Re-ignition of previous trial stimulus during the re-ignition period (300 ms before the second stimulus onset) was accomplished stimulating all excitatory neurons with a non-specific external stimulus. This stimulus increased exponentially with a rate α = 10 s^-1^ as β(1 − *e*^−α(*t*−*t*_0_)^), with β being the reactivation strength and *t*_0_ the time of onset of the stimulus. Reactivation strength was weak (β = 0.17 nA) or strong (β = 2.9 nA).

#### Rate and synaptic tuning

For each simulation in Fig. 3a,b we computed the firing rate (r) and synaptic (s = u · x) tuning, by computing for both measures the difference between neurons within (± 50°) and outside (180° ± 50°) the previous bump location.

