## Supplementary Info for "Interplay between persistent activity and activity-silent dynamics in prefrontal cortex during working memory"

**Supplementary Materials: Supplementary Figures 1-8**

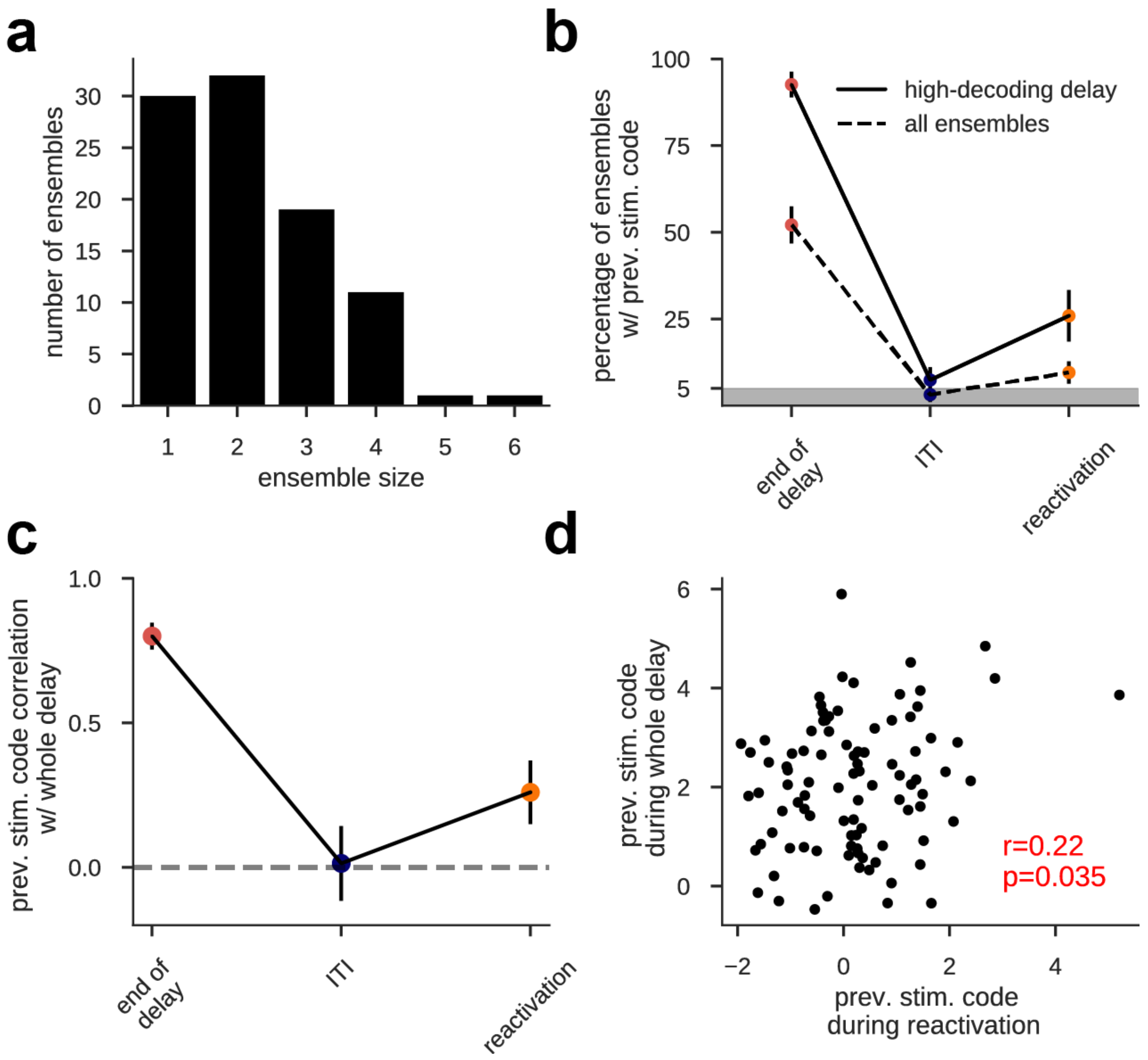


**Supplementary Figure 1. Consistent decoding accuracy in delay and reactivation links these two representations at the neural ensemble level.**

**a)** The size of ensembles of simultaneously recorded neurons varies between 1-6. **b)** Fraction of neural ensembles with significant previous stimulus decoding accuracy ($z > 1.96$, see Methods) computed for all ensembles (dashed line) and only for those ensembles with highest previous stimulus code averaged across the whole delay (see Methods). The incidence of stimulus decoding was significant in delay and reactivation, but not at ITI (binomial test at p=0.05, with n=94 and n=27, for all ensembles and highest delay code, respectively). Error bars are bootstrapped ±s.e.m. **c)** across-ensemble Pearson correlation between delay decoding accuracy (averaged in the whole delay) and decoding accuracy at different time points (p-values: 6.5e-30, 0.87, 0.035). The ensembles with highest delay code also had higher decoding during reactivation, demonstrating the neural association between delay representations and reactivations despite absent code in the ITI. Error bars denote ±s.e.m. computed with a bootstrap procedure. **d)** Individual ensemble values from c, orange.

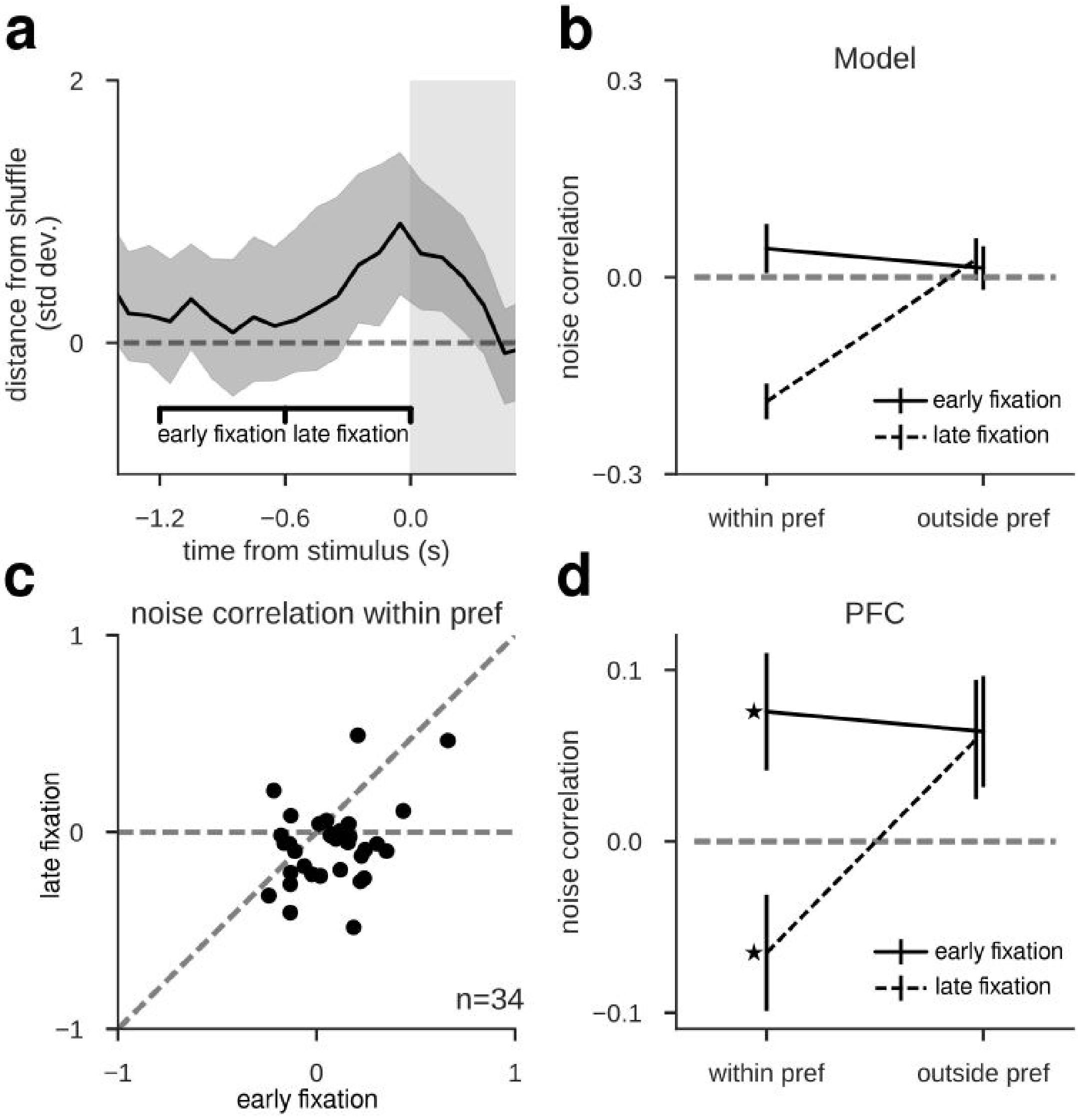


**Supplementary Figure 2. Noise correlation between pairs of neurons is negative at reactivation, as predicted by the model.**

A signature of bump-attractor dynamics is the occurrence of negative pairwise noise correlations for trials where the presented cue lies between the two neurons' preferred locations (*within pref*), but not for other cues (*outside pref*) [6]. As the *within-pref* bump diffuses due to intrinsic noise, it moves closer to one neuron's preferred angle and away from the other neuron's preferred angle, thus increasing one and decreasing the other neuron's firing rate. **a)** Replica of fig. 1c, zooming in on fixation-period to illustrate the periods used in the noise correlation analyses: early fixation with no explicit code for previous-trial cue, and late fixation with code reemergence. Error shading, bootstrapped 95% C.I. . **b)** In the computational model, bump reactivations from subthreshold traces are characterized by negative noise correlations only for trials in the *within-pref* condition and only after reactivation (late fixation), following the onset of the nonspecific input drive (Fig. 3). **c)** In the data, noise correlations for simultaneously recorded PFC pairs selected based on the distance of preferred angles ($60° < \Delta\theta < 120°$) were lower in late fixation than in early fixation for trials in the *within-pref* condition (bootstrap test, p=0.0001, n=34, Cohen's d=0.61). **d)** On average, these PFC pairs were anti-correlated exclusively at the time of reemergent firing rate selectivity in late fixation and *within-pref* cues: Negative noise correlations were specific for late fixation when stimuli were presented *within pref* (ANOVA *trial condition x time point*, F(4)=2.5, p=0.06, n=34). In the *within pref* condition, correlation was negative during late fixation (bootstrap test, p=0.035, Cohen's d=-0.32) but positive during early fixation (bootstrap test, p=0.018, Cohen's d=0.37), and there was a significant difference between late and early fixation (bootstrap test, p=0.0001, Cohen's d=0.61). Moreover, correlations were positive in the *outside pref* condition both during late and early fixation (bootstrap test, p=0.024 and p=0.06, respectively), with no significant difference between late and early fixation (two-sided bootstrap test, p=0.93). In addition, negative noise correlations diminished when using the previous saccade location rather than the previous stimulus as reference (paired bootstrap test, p=0.005, Cohen's d=-0.47), suggesting that the bump diffused only during the delay period, but not after the saccade had been made [6]. Unless stated otherwise, all bootstrap tests were one-tailed in the direction of the model predictions in b). All error bars indicate ±s.e.m.

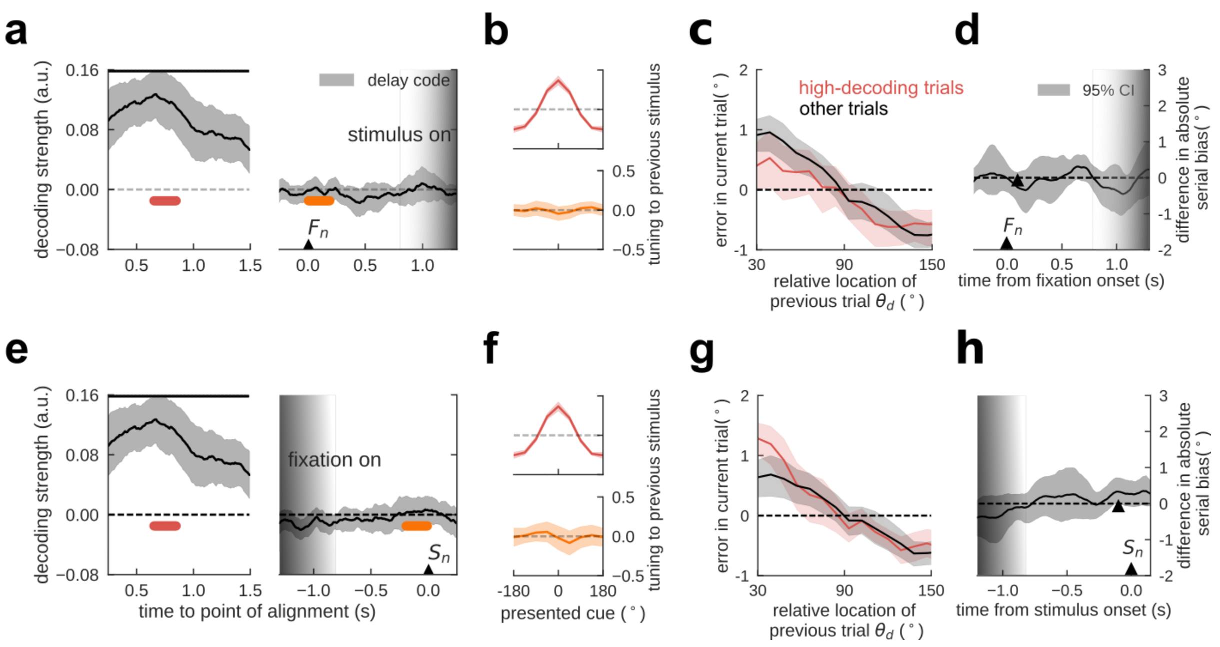


**Supplementary Figure 3. In a dataset with unpredictable stimulus-onset time, previous item representations were not reactivated in the pre-stimulus period.**

We conducted the same analysis as in human EEG (Fig. 2) in a previously published dataset (for experimental details, please refer to the original publication, ref. [35]) with unpredictable fixation period durations (range 0.7 s-1.3 s). Decoding analyses were applied separately for data aligned to the onset of fixation ($F_n$, graded shading indicates range of possible stimulus onset times, upper panels) and aligned to the onset of the stimulus ($S_n$, graded shading indicates possible fixation onset times, lower panels). **a)** Tuning to previous-trial location (decoder trained in delay, .5s **-** 1.0s after stimulus onset) during previous-trial delay (left, stimulus aligned) vanishes in current-trial fixation (right, fixation onset aligned). No reactivation occurs. **b)** Average tuning reconstruction at different epochs for the delay decoder, indicated in a). **c)** Serial dependence separating trials with high (red curve, top quartile) from all other trials' (black curve) decoding accuracy in early fixation (orange in a). Unlike in an experiment with predictable stimulus onset (Fig. 5), serial bias did not differ as a function of decoding strength. **d)** Difference in serial biases (Methods) between *high-decoding* and *other* trials were not significant at any time point in fixation. The black triangle marks the center of 0.2 s decoding window for the split in c. **e-h)** Parallel results were obtained when the analyses of panels a-d were run on data aligned to the time of stimulus onset instead of fixation onset. In d and h, time courses were smoothed using a squared filter of 5 samples. Periods with significant decoding are marked with black horizontal bars, indicating $p<.001$ in a bootstrap test. Shading indicates 95% C.I. in a,d,e and h, and ±s.e.m. in panels c and g.

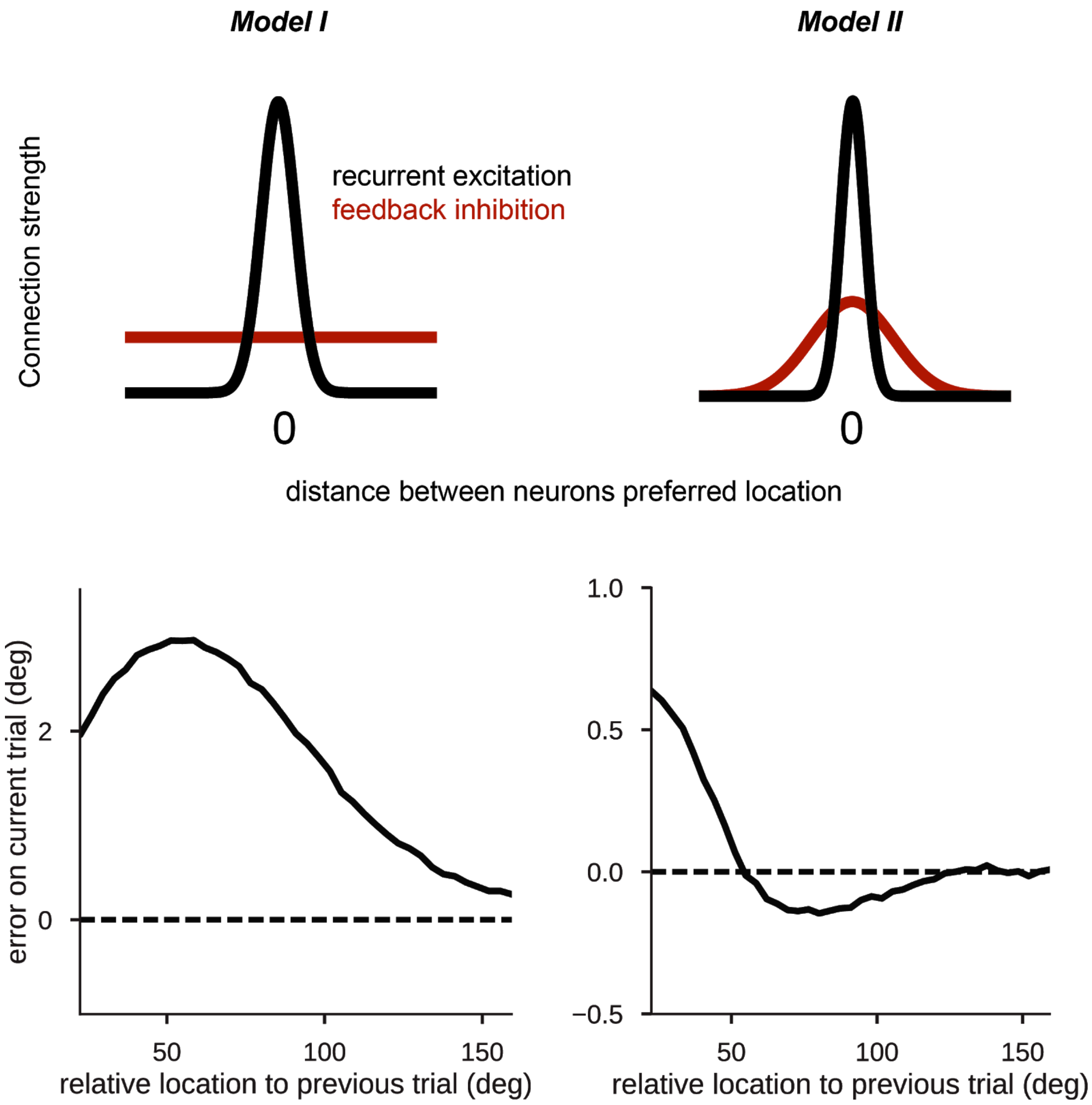


**Supplementary Figure 4.** Structured inhibition is necessary for repulsive serial biases at far distances. Top panel, illustration of two different models that have different inhibitory connectivity profiles. On the left, inhibitory connectivity strength from inhibitory to excitatory neurons is similar for all distances between their preferred locations. On the right, inhibition is structured such that similarly tuned neurons have stronger feedback inhibition.

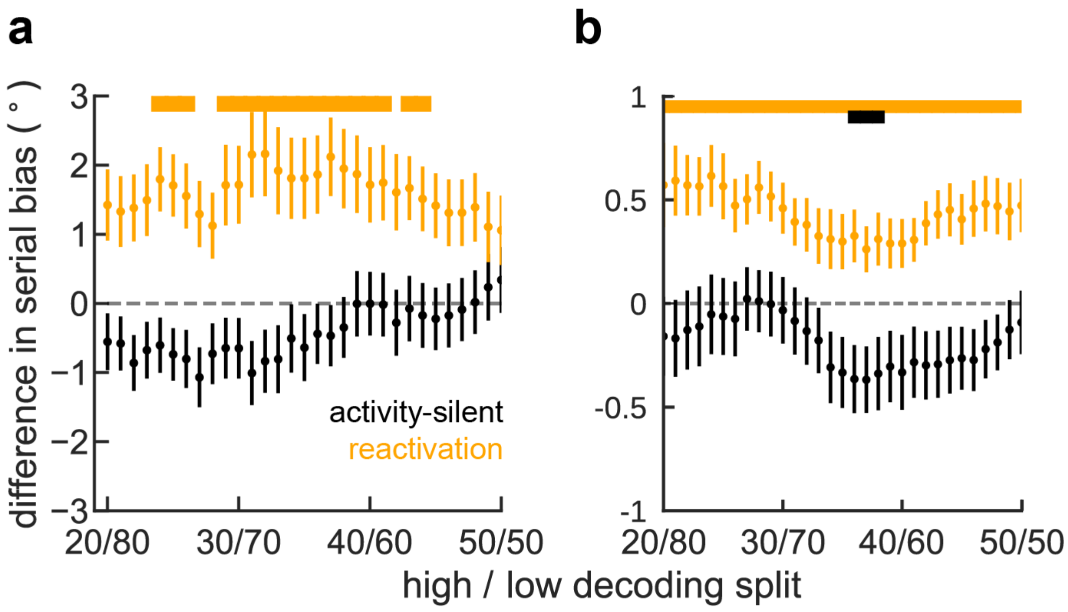


**Supplementary Figure 5. The more trials are included in the split between high- and low-decoding trials, the lower the serial bias**

**a)** In monkey behavior **b)** In human behavior. Error bars are ±s.e.m. and colored bars mark where corresponding difference in serial biases is different than zero ($p<0.05$, bootstrap).

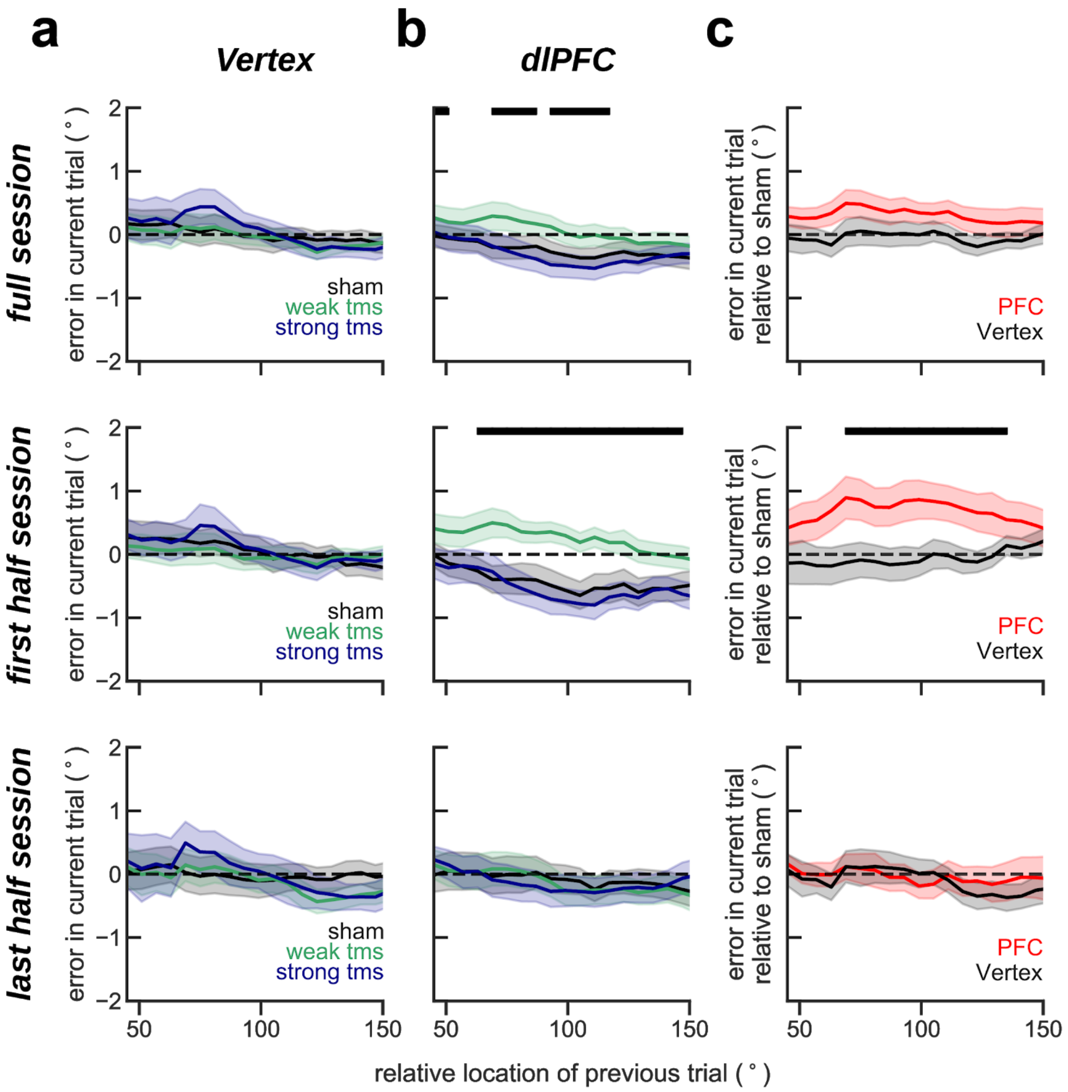


**Supplementary Figure 6. The effect on serial biases of targeting dlPFC with TMS diminishes in the course of the experimental session.**

Same analyses as in Figure 6, but (top) analyzing trials from the full session, (middle) first half session (225 trials, replication of Figure 6) and (bottom) last half session (225 trials). The behavioral impact of PFC TMS stimulation declined through the session, as if subjects desensitized (*prev-curr* ⨯ *TMS intensity* ⨯ *session-half* $t_{11083}$=-2.38, p=0.017. Methods, *Linear Mixed Models*). Serial biases were modulated by TMS in PFC, but not in Vertex (*prev-curr* ⨯ *TMS intensity* ⨯ *coil location*, $t_{18272}$=2.21, p=0.027. For dlPFC: *prev-curr* ⨯ *TMS intensity*, $t_{11087}$=2.13, p=0.032. For Vertex: $t_{7166}$=0.03, p=0.97. Methods, Linear mixed models) when analyzing the full session, and analyzing only the first half session ($t_{9133}$=2.51, p=0.011).

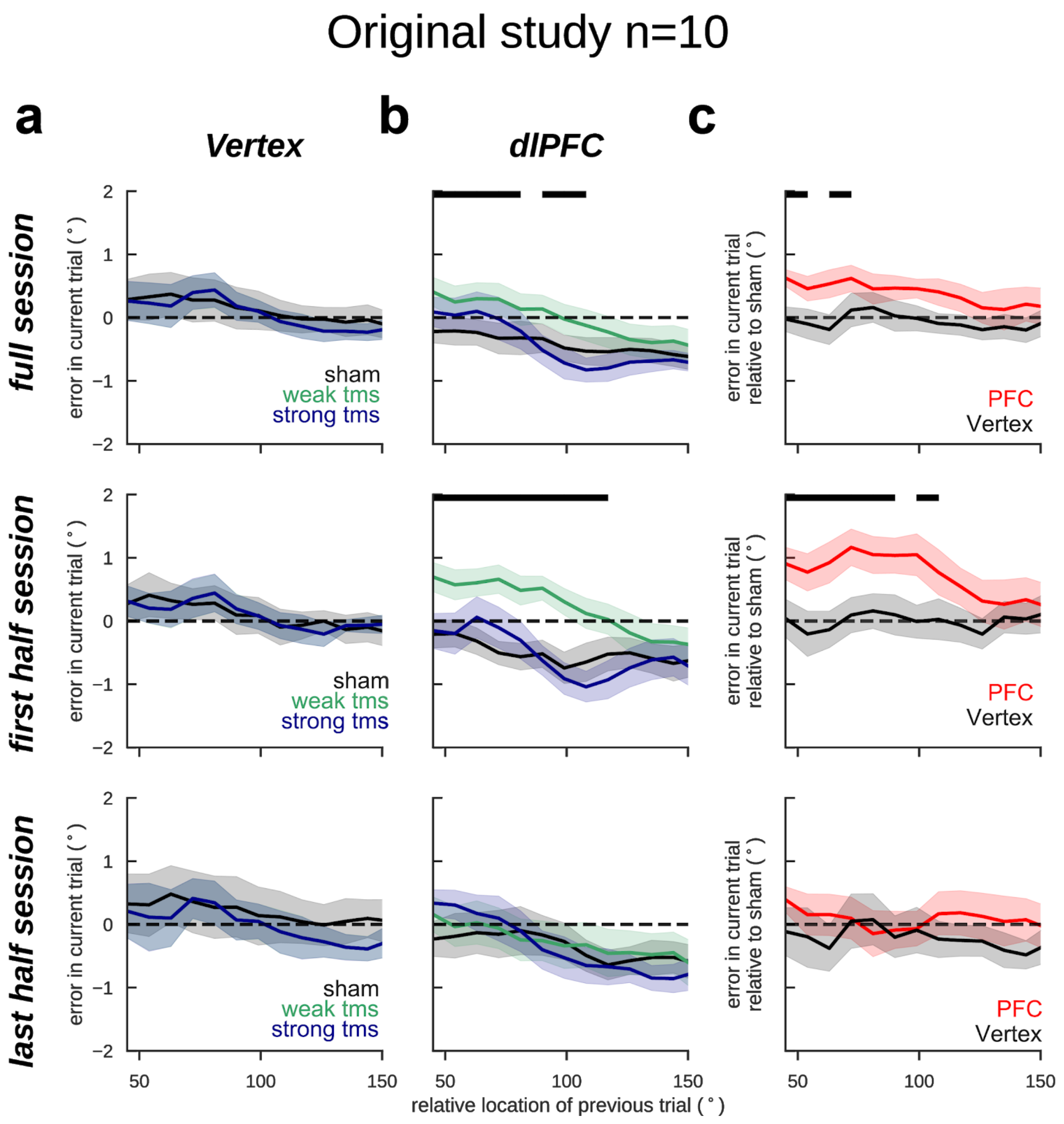


**Supplementary Figure 7.** Same as Supplementary Fig. 6, but only analyzing data from the original study (n=10). Similarly to when pooling both the original and replication studies together, the behavioral impact of PFC TMS stimulation declined throughout the session, however not significantly (*prev-curr* × *TMS intensity* × *session-half* $t_{5701}$=-1.73, p=0.08. Methods, Linear Mixed Models). Serial biases were modulated by TMS in PFC, but not in Vertex ($t_{5705}$=1.92, p=0.05) when analyzing the full session, and analyzing only first half session ($t_{3059}$=2.59, p=0.009, Methods).

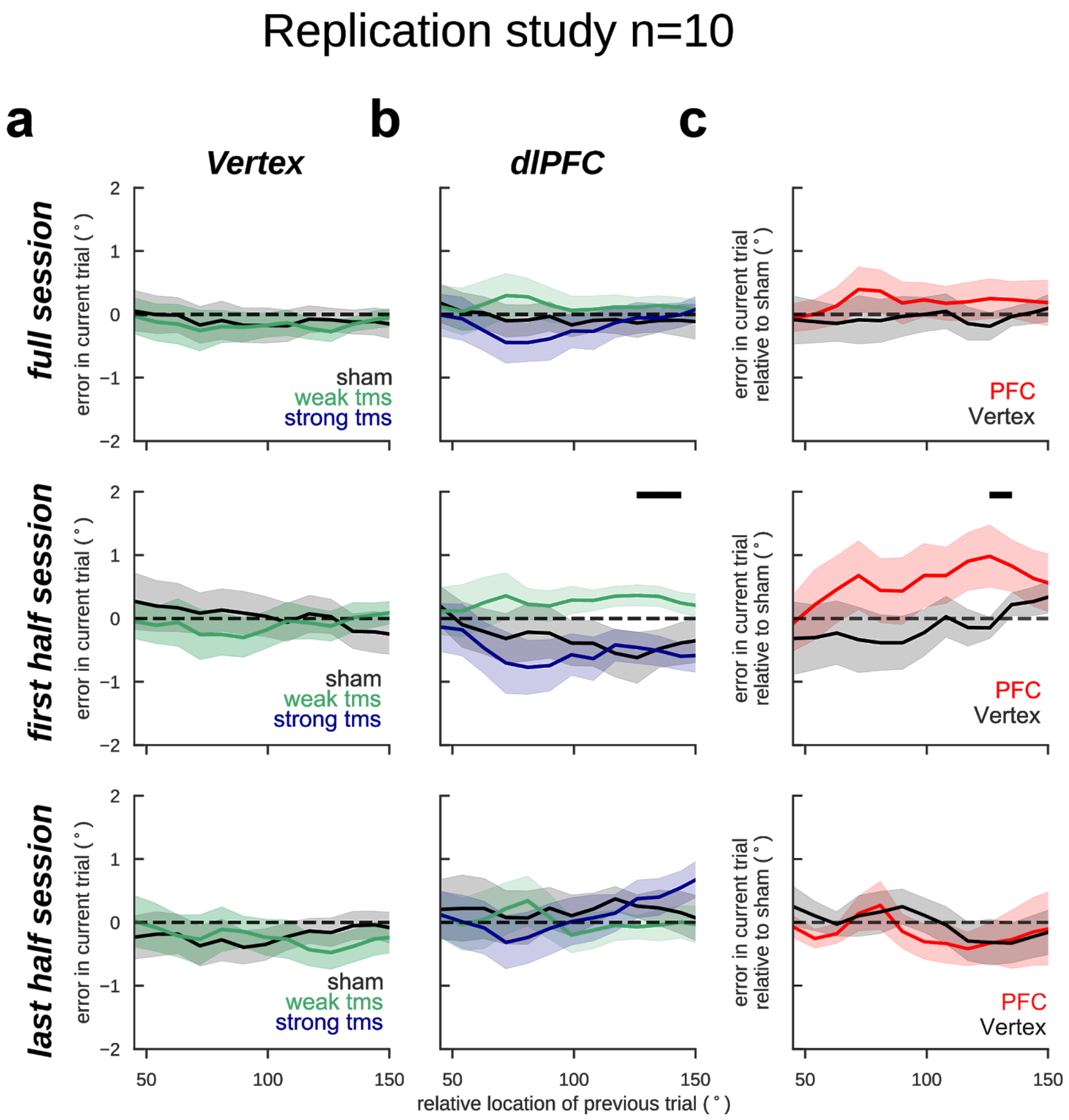


**Supplementary Figure 8.** Same as Supplementary Fig. 6 and 7, but only analyzing data from the pre-registered (https://osf.io/rguzn/) replication study (n=10). Similarly to the original experiment, the behavioral impact of PFC TMS stimulation declined throughout the session, however not significantly (*prev-curr* × *TMS intensity* × *session-half* $t_{5375}$=-1.63, p=0.1. Methods, Linear Mixed Models). Similarly to the original study, serial biases were more strongly modulated by TMS in PFC than in Vertex, however not significantly ($t_{5379}$=1.12, p=0.25) when analyzing the full session and the effect was stronger when analyzing only the first half-session ($t_{2675}$=1.91, p=0.06, Methods).

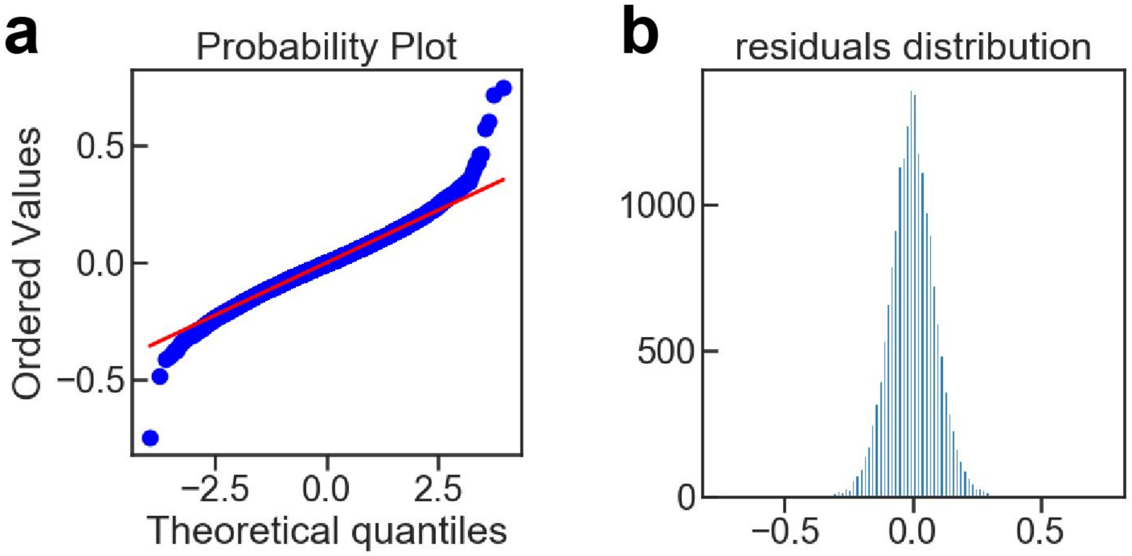


**Supplementary Figure 9.** qqplot (a) and distribution (b) of residuals for the linear mixed model applied in the TMS data analysis (Fig. 6).
